# SIRT1 regulates DNA damage signaling through the PP4 phosphatase complex

**DOI:** 10.1101/2022.09.29.510108

**Authors:** George Rasti, Maximilian Becker, Berta N. Vazquez, Maria Espinosa-Alcantud, Irene Fernández-Duran, Andrés Gámez-García, Jessica Gonzalez-Nieto, Laia Bosch-Presegué, Anna Marazuela-Duque, Sandra Segura-Bayona, Alessandro Ianni, Joan-Josep Bech-Serra, Michael Scher, Lourdes Serrano, Uma Shankavaram, Hediye Erdjument-Bromage, Paul Tempst, Danny Reinberg, Mireia Olivella, Travis Stracker, Carolina de la Torre, Alejandro Vaquero

**Affiliations:** Chromatin Biology Laboratory, Josep Carreras Leukaemia Research Institute (IJC), Ctra de Can Ruti, Camí de les Escoles s/n, 08916, Badalona, Barcelona, Spain; Chromatin Biology Laboratory, Cancer Epigenetics and Biology Program (PEBC), Bellvitge Biomedical Research Institute (IDIBELL), Av. Gran Via de l’Hospitalet, 199-203, 08908 L’Hospitalet de Llobregat, Barcelona, Spain; Tissue Repair and Regeneration Lab, Sciences and Methodology Department, Universitat de Vic, Universitat Central de Catalunya, Vic, Spain; Institute for Research in Biomedicine (IRB Barcelona), The Barcelona Institute of Science and Technology, Barcelona, Spain; The Francis Crick Institute, 1 Midland Road, London NW1 1AT, UK; Department of Cardiac Development and Remodeling, Max-Planck-Institute for Heart and Lung Research, Ludwigstrasse 43, 61231 Bad Nauheim, Germany; Proteomic Unit, Josep Carreras Leukaemia Research Institute (IJC), Ctra de Can Ruti, Camí de les Escoles s/n, 08916, Badalona, Barcelona, Spain; Howard Hughes Medical Institute, Division of Nucleic Acids Enzymology, Department of Biochemistry, University of Medicine and Dentistry of New Jersey, Robert Wood Johnson Medical School, New Jersey 08854 USA; Department of Science, BMCC, The City University of New York (CUNY), 199 Chambers Street N699P, New York, NY 10007, USA; Radiation Oncology Branch, National Cancer Institute, Bethesda, Maryland, 20892, USA; Molecular Biology Program, Memorial Sloan-Kettering Cancer Center, New York, New York 10065, USA; Department of Cell Biology, New York University School of Medicine, New York, NY 10016, USA; Howard Hughes Medical Institute, Department of Biochemistry, New York University School of Medicine, New York, 10016, USA; Bioinfomatics and Medical Statistics Group, Faculty of Science, Technology and Engineering. University of Vic-Central University of Catalonia, Vic, Spain

## Abstract

The Sirtuin family of NAD^+^-dependent enzymes plays an important role in maintaining genome stability upon stress. Several mammalian Sirtuins have been linked directly or indirectly to the regulation of DNA damage during replication through Homologous recombination (HR). The role of one of them, SIRT1, is intriguing as it seems to have a general regulatory role in the DNA damage response (DDR) that has not yet been addressed. SIRT1-deficient cells show impaired DDR reflected in a decrease in repair capacity, increased genome instability and decreased levels of γH2AX. Here we unveil a close functional antagonism between SIRT1 and the PP4 phosphatase multiprotein complex in the regulation of the DDR. Upon DNA damage, SIRT1 interacts specifically with the catalytical subunit PP4c and promotes its inhibition by deacetylating the WH1 domain of the regulatory subunits PP4R3α/β. This in turn regulates γH2AX and RPA2 phosphorylation, two key events in the signaling of DNA damage and repair by HR. We propose a mechanism whereby during stress, SIRT1 signaling ensures a global control of DNA damage signaling through PP4.

## INTRODUCTION

The members of the Sirtuin family of NAD^+^-dependent enzymes play a key role in the response to metabolic, oxidative and genotoxic stress(1). Sirtuins coordinate an efficient adaptation to these stress conditions at different levels including protection of genome stability, metabolic homeostasis, and increased survival. Among the most relevant roles of Sirtuins is their direct implication in maintaining genome integrity which involves chromatin organization and structure, mitotic progression and DNA repair(2,3). In this functional context, Sirtuins have been implicated in the regulation of both DNA damage response (DDR) signaling and DNA repair itself(4).

The DDR is tightly regulated by a wide range of protein post-translation modifications. Among the most relevant and conserved of them is phosphorylation of histone and non-histone proteins, which participate in all crucial steps of DDR. Phosphorylation of serine 139 in the histone H2A variant H2AX (γH2AX) is a hallmark of double strand break (DSB) signaling in the DDR(5–7).The establishment of γH2AX, catalyzed by the kinases ATM, ATR or DNA-PKcs is an early step in the DDR that promotes interactions and sequential recruitment of other key regulatory factors such as the MRN complex (MRE11-RAD50-NBS1) and MDC1(8,9). After DNA repair is completed, γH2AX is dephosphorylated allowing the cell to deactivate the DNA damage checkpoint and to resume cell cycle(10). Several phosphatases have been described to play this role upon completion of repair including PP2A, PP4 or Wip1(11). Among them, PP4 it was shown to antagonize ATR signaling during replication stress, and its depletion induces γH2AX hyperphosphorylation and a block in DNA damage efficiency(12,13).

PP4 phosphatase is a Ser/Thr PP2A-like multisubunit complex involved in many stress-associated processes beyond the DDR, such as NF-kB signaling, immune response, meiosis, centrosome maturation, splicing, cell cycle regulation and glucose metabolism(14). Reflecting its physiological relevance, PP4 is frequently deregulated in a wide range of cancers(15,16). The PP4 complex is a trimeric/tetrameric complex containing a catalytic subunit PP4c and different combinations of the regulatory subunits PP4R1, PP4R2, PP4R3α and PP4R3β (also known as SMEK1 and 2, respectively), and PP4R4(17). Given the high degree of identity between PP4c and the PP2 catalytic subunits (65%), it is the regulatory subunits that mediate the specific roles of PP4, including binding to different substrates and other specific factors(18,19). Despite the important role of this complex, little is known about the way this activity is regulated. The most clearly conserved PP4 complex is formed by PP4c-R2-R3 (3α and 3β in mammals), that is responsible for γH2AX dephosphorylation and hypersensitivity to DNA damage-inducing agents, like cisplatin^59^. PP4R2 is required to stabilize PP4c structure and its deficiency decreases the specific activity of PP4c(12,20). In contrast, PP4R3α and β play a more complex role. While they are key to recognize substrates through their N-terminal WH1 domain, they also seem to restrict PP4c activity *in vitro*, suggesting that they control PP4 enzymatic activity(21). Besides γH2AX, the PP4c-R2-R3 complex also dephosphorylates other key targets such as RPA2, KAP1 or 53BP1 in the context of DDR (22,23). RPA2 is a subunit of the trimeric RPA complex, which binds to ssDNA in the repair of DSBs and base damage. RPA2 phosphorylation by ATR on S33 and T21 during the S and G2 phases(60) participate in HR pathway regulation, replication stress tolerance and mitotic exit, among other functions(24). Thus, RPA2 phosphorylation is associated with DSB pathway choice, as RPA2 dephosphorylation induces foci formation of the HR-specific factor RAD51(25).

So far, Sirtuins SIRT1,2,6 and 7 have been linked to the DDR. Of them, SIRT1 and SIRT6 seem to have a more prominent roles, and in contrast to SIRT2 or SIRT7, their loss impairs the phosphorylation of H2AX, which results in reduced DNA repair efficiency(26–28). SIRT6 has a major role in the initial steps of DSB repair as a pioneering factor that recognizes and binds to DNA damage, initiating the DSB repair process(29). SIRT6 loss impairs recruitment of HR and non-homologous end-joining (NHEJ)-associated factors, including MRE11, BRCA1, 53BP1, Ku80 and results in decreased levels of γH2AX(29). SIRT1 loss inhibits foci formation of NBS1, RAD51, BRCA1 and γH2AX(30,31). However, although several SIRT1-dependent mechanisms have been described involving deacetylation of NBS1, Ku70, WRN, MOF or TIP60, the underlying mechanism and how are they interconnected in the context of a global SIRT1 function, are not well understood(61). The most paradoxical case is γH2AX as deacetylation of NBS1 by SIRT1 or the reported relationship between SIRT1 and ATM cannot explain the observed decrease of this modification(32–34). Here we identify and characterize a direct functional antagonism between SIRT1 and PP4 complex through deacetylation of regulatory subunits PP4R3α and PP4R3β. Our evidence helps to explain the decreased γH2AX levels observed in *Sirt1^−/−^* cells and suggests a global coordinated control of DDR progression by SIRT1 through regulation of PP4 function.

## MATERIALS AND METHODS

### Reagents

camptothecin (CPT) and hydroxyurea(HU) were obtained from Sigma-Aldrich, okadaic acid(OA) from Calbiochem and Puromycin from Invivogen. The following antibodies were used: HA (Sigma-Aldrich, H6908), FLAG (Sigma-Aldrich F1804), c-Myc (Cell Signaling2276S), actin (Sigma-Aldrich A1978), α-tubulin (Sigma-Aldrich T6199), SIRT1 (Millipore, 07-131) SIRT6 (ABCAM AB62739), histone H3 (Cell Signaling, 9715), GFP (MERCK, MAB2510), RPA32 (Cell Signaling 2208), PP4R2 (Bethyl, A300-838A), PP4R3α (Bethyl, A300-840A), PP4R3β (Bethyl, A300-842A), PP4C (Abcam, ab171870), Anti-phospho-Histone H2A.X (Ser139)(abcam ab 2893 and Merck Millipore, JBW301), phospho-RPA2 (Ser33) (Novus, NB100-544 and Bethyl, A300-246A), phospho-RPA2 (Ser4/8) (Sigma, PLA0071).

### Cell culture studies

SIRT1 wild type and SIRT1−/− Mouse embryonic fibroblasts (MEFs) were cultured in DMEM supplemented with 20% fetal bovine serum, Pen-strep (10000 IU/mL, 10 mg/mL), non-essential amino acids and sodium pyruvate according to the manufacturer’s instructions. Hela and 293T cells (Human Embryonic Kidney, HEK293T) were cultured in Dulbecco’s modified Eagle’s medium (DMEM) (GIBCO, Invitrogen, Carlsbad, CA, USA) supplemented with 10% fetal bovine serum. All the Cells were grown at 37°C in an atmosphere containing 5% CO2, and 100% humidity. At 48 hours of transfection (If indicated) the cells were treated with the following conditions before harvesting at indicated times: 2mM of H2O2 (MERCK) for 1 hour, 5mM hydroxyurea (Sigma-Aldrich) for 4 hours, 1 μM campthotecin (Sigma-Aldrich) for 1 hour and irradiation with X rays at 7.5 and 10 Gy. The cells were exposed to different concentrations of EX527(1 and 10 μM) during 24 hours.

### Gel filtration analysis

SIRT1 was purified from 2 grams of HeLa nuclear extract and loaded on a phosphocellulose P11 cation exchange column (Whatman, USA) in a BC buffer (50mM Tris-HCl pH 7.9, 0.2mM EDTA, 10% Glycerol, 1mM DTT, 0.2mM PMSF and the indicated concentration of KCl) with a gradient from 100mM to 1M KCl. The Flow-Through (FT) of the P11 column was loaded on the anion exchange column DE52 and resolved in a linear gradient of BC buffer from 100mM to 600mM KCl. The SIRT1-containing fractions were pooled and dialyzed to BC buffer 80mM KCl and fractionated on the weak anion exchange column DEAE-5PW (Diethylaminoethyl cellulose 52; TosoHaas, Montgomeryville, USA). The DEAE column was resolved by using a linear gradient of 80mM to 725mM KCl in buffer BC. The fractions containing SIRT1 were precipitated with saturated Ammonium sulphate 5M Tris 50mM pH7.9 at 40%. The precipitate was then resuspended in BD buffer (BC buffer with 50mM Ammonium sulphate) with 500mM KCl and fractionated by molecular weight on the gel filtration column Sephacryl S-400 (GE healthcare, USA) with a fractionation range of 2×104–8×106 Da. Purified SIRT1 was then dialyzed to BD buffer 1M KCl, applied to hydrophobic interaction chromatography on a Phenyl-Superose column and resolved in a linear gradient of BD buffer from 1M to 350 mM KCl. SIRT1 eluted around BD buffer 700mM KCl and the fractions were pooled and dialyzed to BC buffer with 40mM KCl. Purified SIRT1 was subjected to a MonoQ anion exchange column (GE healthcare, USA) and resolved in a linear gradient from BC buffer with 40mM to 400mM KCl. The Purified fractions containing SIRT1 were subjected to SDS-PAGE and visualized by silver staining. The fractions 25 and 26 were pooled, loaded on a single lane and subjected to SDS-PAGE followed by staining with colloidal blue (Invitrogen, USA). Bands were cut and containing proteins were identified by mass-spectrometry ((see Supplementary information).

### Viral generation and infections

To produce retrovirus, Platinum A cells were seeded into culture dishes a day before transfection in DMEM supplemented with 10% FBS. The cells were co-transfected with 2 μg of pVSVG plasmid and 8 μg of the retroviral vector and supernatant media was collected after 24h and 48h and stored at −80°C. For the infection, cells were plated 24h before infection in 6-well plates. Then, media was replaced by the stocked supernatant containing the retrovirus and incubated for 16 hours, thereafter the media was replaced by fresh media and after 24 hours cells were selected with puromycin. For shRNA knockdown with lentiviral infection, Mission shRNAPP4C (pLKO.1-puro, Clon ID #TRCN0000080835) were obtained from Sigma-Aldrich, Inc. HEK293T cells were co-transfected with pMD2.G and pPAX2 (Addgene) together with pLKO-shRNA constructs (PLKO.1 shRNA PP4C) or a non-targeting shRNA (pLKO.1-puro ID # SHC003). Viral supernatants were harvested after 48 h. For infections, SIRT1 MEF cells were incubated with viral supernatants in the presence of 8 μg /ml polybrene. After 48 h, puromycin (5μg /ml) as selection was added to the medium. Cells were grown in presence of selection for 4-6 days to select for stable knockdown cells.

### Invitro enzymatic reactions

For the phosphatase assays, PP4 complex subunits were overexpressed in HeLa cells or SIRT1 MEFs and purified using Flag resin to immunoprecipitate the catalytic subunit PP4C. The bound proteins were eluted by competition with a large excess of free FLAG peptide (0.4 mg/mL). In vitro Phosphatase assays were performed in Phosphatase Reaction Buffer (PRB, 50 mM Tris pH 7.2, 0.1 mM CaCl2, 5 mM MnCl2 and 0.2 mg/ml BSA). Purified proteins were first equilibrated with PRB for 10min at 37°C followed by addition of phospho-H2AX-enriched histone as a substrate and incubation at 37°C for 30min and analyzed by Western-blot. For the deacetylation assays HA-SIRT1 and PP4 complex subunits were expressed separately in Hela cells, immunoprecipitated and purified using HA and Flag peptides in BC100 buffer. Deacetylation assays were performed in deacetylation buffer (50 mM Tris HCl pH 8, 100mM NaCl, 2mM DTT, 5% Glycerol) with or without 5 mM of NAD+. SIRT1 was first equilibrated with deacetylation buffer for 10min at 37°C. Then, hyperacetylated histone substrate were added to the reaction mixture and incubated at 37°C for 1h and analyzed by Western-blot.

In the case of PP4R3α/β deacetylation by SIRT1, Biotinylated peptides (Biotin-PNTAYQK(Ac)QQDTLI) previously immobilized on magnetic Streptavidin beads (Dynabeads MyOne Invitrogen™ 65601) were incubated in presence or absence of SIRT1 −/+ NAD+ (0.5 nM) in a total volume of 100 μl of deacetylase buffer (60 mM Tris HCl pH 7,8, 40 mM MgCl2, at pH 7.0, 2mM DTT and protease inhibitor cocktail).The reaction mixtures were incubated at 37 °C for 90 minutes and the beads were washed three times with BC100 and deacetylation of Ac-PP4R3A/B (K64) by SIRT1 was detected by MALDI-TOF (see Supplementary information).

### Analysis of metaphase Aberrations

Cells were treated with 250nM Mitomycin C for 24h and then arrested in metaphase with 100 ng/ml of Colcemid for 3 hour and fixed in 3:1 methanol:acetic acid. Resuspended cells were dropped onto a slide and stained with Giemsa stain (1:12 in 3% methanol), and they were allowed to rock for 20 minutes. They were rinsed with water in a Coplin jar for 5 minutes while shaking. Images were acquired with an 100× obejctive(100×) and Metafer software (MetaSystems). Metaphases were analyzed using ImageJ.

### HR functional assays

The SceI-GFP reporter was used to measure HR repair as described elsewhere (64). Briefly, U2OS-HR-DR cells were transfected with SCR, SIRT1 and/or PP4C siRNA for 24h using Dharmafect reagent. To induce I-SCEI DSB, cells were incubated with 1μM Shield- 1 and 100nM triamcinolone acetonide for an additional 48h, moment at which cells were trypsinized and analyzed for GFP expression in a FACS canto cytometer. Data was analyzed using Flowjo and results were expressed as the mean percentage of GFP+ cells from three independent experiments.

### Cell cycle analysis

Cell cycle analysis was performed using the Click-iT EdU Alexa Fluor 647 Flow Cytometry Assay kit (C10340, ThermoFisher Scientific). Cells were treated with 10 mM EdU and incubated for 30 min before being trypsinized. After washing with PBS, cells were fixed with 4% PFA for 15 min, washed once with washing buffer (1% BSA, 2 mM EDTA in 1× PBS) and permeabilized (0,5% Triton, 1% BSA in 1× PBS) for 10 min. After washing with washing buffer, click chemistry using Alexa Flour 647 azide was performed for 30 min in the dark, as stated in the manufacturer’s instructions. Cells were then washed again in washing buffer and resuspended in analysis buffer (0,02% (wt/vol) sodium azide, 0.5 mg/mL DAPI in 1 mg/ml BSA in 1× PBS). Cell cycle was analysed using a BD FACSCanto™ II System and the FlowJo 7.6 software. Standard gating for cells versus debris and singlet was conducted.

### Proliferation and viability

Cell proliferation and viability were measured at the indicated time-points using the trypan blue exclusion assay. Briefly, supernatant and attached cells were collected together, centrifuged 5 min at 1500 rpm, and resuspended in complete media. 20 mL of cells were diluted in 40 ml of trypan blue. 10 ml sample were loaded on to a hemocytometer, and total, alive and dead cells were counted using a Leica DMi1 inverted microscope.

### TCGA proteomics, phosphoproteomics and transcriptomics

To evaluate correlations between SIRT1, SIRT6, PPP4C protein abundance and pRPA2 (p238), pTRIM28 (pS473) and pATM (pS1981) levels in the Breast Invasive Carcinoma (TCGA) dataset, mass spectrometry levels measured by the National Cancer Institute’s Clinical Proteomic Tumor Analysis Consortium (CPTAC) were accessed and downloaded through the cBioportal web-based utility (http://www.cbioportal.org). Pearson correlations were calculated using GraphPad Prism version 8.0. A subset of patients with mutations or deep deletions in HDR genes was defined (see Figure 7D) and gene expression and protein levels correlation in this subset was conducted.

### Neutral comet assay

To detect DSBs, the neutral comet assay was performed as described previously (65). U20S and SIRT1 MEFs Cells were infected with either control or PP4c-specific siRNA and treated or not with camptothecin 1μM for 90 minutes. The images were acquired using a Leica AF-5000 confocal microscope and analyzed and quantified using the Comet Score program.

### FRET

FRET assays were performed as previously described^66^. Human SIRT1 and PP4C were amplified by PCR with Sal-I and BamH-I sites added to the 5’ and 3’ ends, respectively. The PCR products were cloned into the mCerulean-C1 and YFP-C1 vector so that both proteins were either N-terminally tagged with the chromophore. All constructs were confirmed by DNA sequencing. HeLa cells were co-transfected with 8μg YFP-SIRT1 and 1μg CFP-PP4C plasmid DNA per plate and plated on a coverslip. After 48h, transfected cells were imaged using a Leica TCS SP5 confocal imaging system. To measure FRET signals by acceptor photobleaching, we imaged the fluorescent intensity in a selected region of interest. YFP-SIRT1 fluorescence was then photo-destroyed by repeated scanning with the 568-nm laser line. The fluorescence intensities of images of pre-bleached CFP-PP4C were compared to corresponding regions of post-bleached images. As controls, non-fused chromophores were paired with their corresponding protein-fused chromophores and similarly processed. FRET was measured as an increase in CFP fluorescence intensity following YFP photobleaching, and calculated as 100 × [(CFP post-bleach – CFP pre-bleach)/YFP post-bleach]; taking into account YFP and CFP background noise.

### Phospho-H2AX-enriched Histone extraction

Hela cells were treated with 2mM of H2O2 (MERCK) for 1 hour, then the cells were scraped down by a cell scraper and washed with PBS and pelleted by centrifugation at 4000 rpm for 1 minute, subsequently the cells were resuspended in buffer A (10 mM Tris, pH 7.9, 1,5 mM MgCl2, 10 mM KCl, 0,1 mM PMSF, 0,5 mM DTT, supplemented with Protease Inhibitor(sigma)) for 10 minutes on ice, centrifuged at 12000 × g for 1 minute. The pellet (nuclear fraction) was resuspended with 0.5 M HCl (500 μl/ 10 cm2 plate). The samples were resuspended by vortex and then incubated on ice for 15 minutes. After centrifugation at 14000 × g for 10 min at 4°C, the supernatant (soluble acid proteins) was collected and precipitated with TCA in final concentration of 20% and incubate on ice for 1 hour. After centrifugation at 14.000 × g for 10 min at 4°C, the pellet was washed with 100% ice-cold acetone and then centrifuged at 14000 × g for 10 min at 4°C. After drying the pellet in RT, it was resuspended with BC100 and stored at −80°C.

### Statistical analysis

The represented values show means of at least three of more independent experiments (n≥3) with error bars representing standard error of mean (SEM) unless otherwise specified. Unless stated otherwise, statistical analysis of the data was performed using two-tailed Student’s t test (α = 0.05).

Further Materials and Methods information can be found in supplementary information file.

## RESULTS

### Purification of endogenous SIRT1 reveals a direct interaction with the PP4 complex

To further understand the role of SIRT1 in genome stability, we purified endogenous SIRT1 to identify any interacting proteins that may shed light on its role in the DDR. Nuclear extracts from HeLaS3 were processed through a sequence of chromatography steps (Figure 1A). Following the final step of an anion exchange monoQ column, proteins were run in SDS-PAGE gel and silver-staining revealed co-elution of the SIRT1 band with three different protein bands of lower MW (Figure 1B). Mass spectrometry analysis identified the members of a previously described PP4 complex, the catalytic subunit PP4c and regulatory proteins PP4R2, PP4R3α and PP4R3β (Figure 1C)(12). Given the well-established role of this PP4c-R2-R3α-R3β complex in DNA damage response and repair(12), this finding suggested that SIRT1 could participate in the role of PP4c in DNA repair.

**Figure 1:**
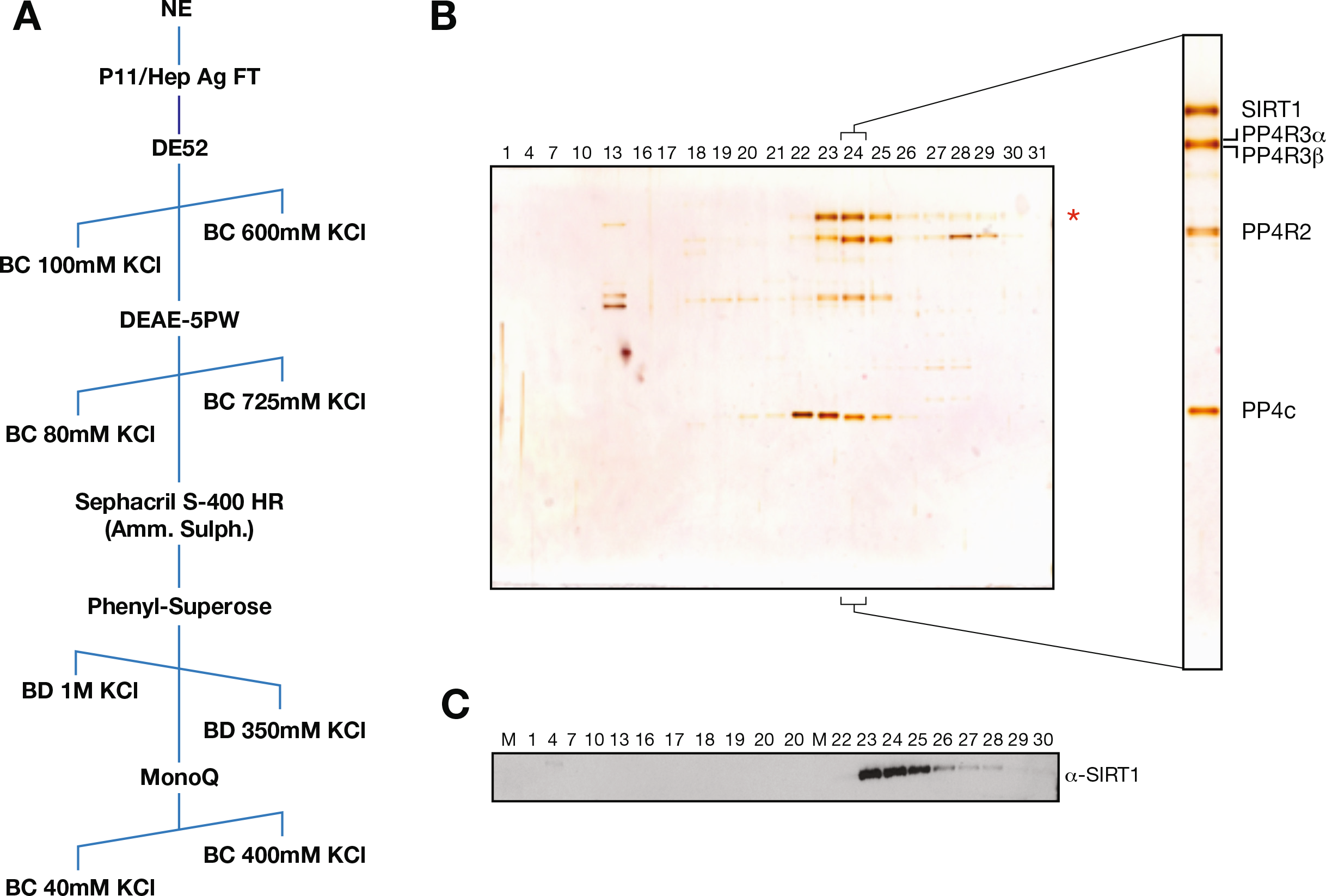
Identification of *a novel protein complex containing SIRT1 and the PP4 phosphatase complex*. **(A)** Schematic representation of the SIRT1 purification from HeLaS3 nuclear extracts (NE) including the buffers (BC or BD) and range of KCl concentrations (from 40mM to 1M) in each step (see Methods and Supplementary information). **(B)-(C)** Silver stained SDS-egl (**B**) and western-blot **(C)** of the anion-exchange MonoQ column, the final chromatography step of the purification. Fractions 23-25 were pooled and analyzed by mass spectrometry. The analysis showed that PP4 complex components PP4c, PP4R2, PP4R3α and PP4R3β co-fractionate with SIRT1 (Table S1).

### SIRT1 interacts with the PP4 complex upon genotoxic stress

Given the functional relationship between both factors and the DDR, we tested whether the interaction between SIRT1 and the PP4 complex was specifically boosted by specific genotoxic stress conditions. Surprisingly, we observed that although the interaction took place under all tested conditions, including replicative stress (HU and CPT) and ionizing irradiation (IR), it was specifically promoted upon oxidative stress (Figure 2A and Supplementary Figure S1A). This oxidative stress-dependent interaction was also observed *in vivo* using FRET assays between Cerulean (CFP)-PP4 and YFP-SIRT1 (Figure 2B and Supplementary Figure S1B). We further confirmed that stress conditions drive the formation of a stable SIRT1/PP4 complex, as we detected the cofractionation of SIRT1 and PP4 subunits in an approx. 600kDa complex in gel filtration analysis (Figure2C), corresponding roughly to the sum of the homotetramer SIRT1 complex(35) and the PP4c complex (400 and 240 kDa, respectively) (36).

**Figure 2:**
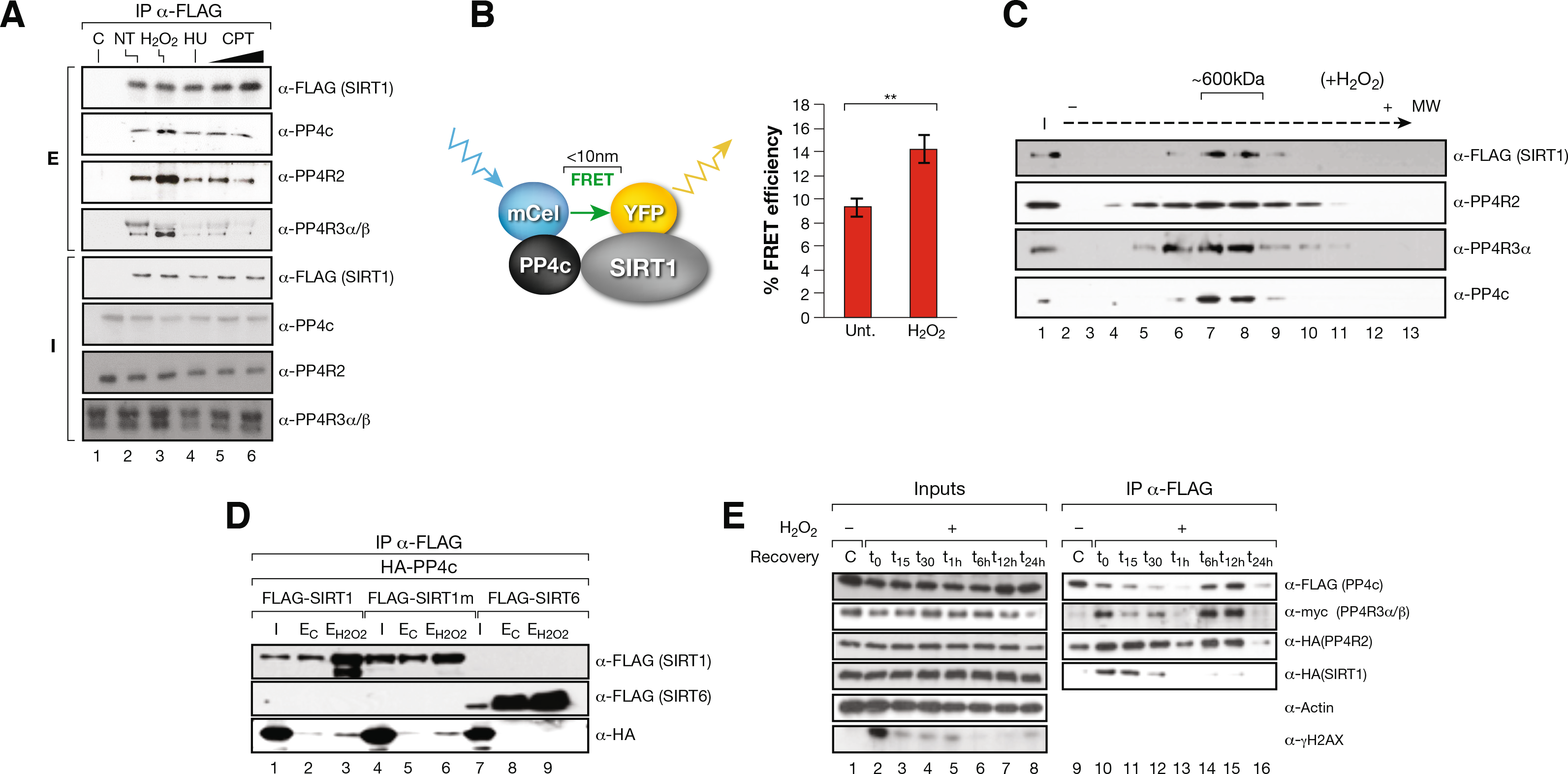
SIRT1-PP4 complex formation in boosted by oxidative stress conditions. **(A)** SIRT1 and the PP4 complex interaction with SIRT1-FLAGPI experiments in HeLa cells non-stress (NT) or under 5mM H_2_O_2_ (2h), 2mM hydroxyurea (4h, HU) and 10 and 20 μM camptothecin (2h, CPT). **(B)** FRET analysis of SIRT1-YFP (acceptor) and PP4C-CFP (donor) in Hela cells treated during 1h with 2mM H_2_O_2_. Cells were imaged with confocal fluorescence microscopy and fluorescence of CFP-PP4C was measured before and after photobleaching in control and oxidative stress conditions. Left, schematic representation of the process. Right, uqantification of detected FRET efficiency is shown from n=3 experiments. Two-tailed T-test analysis (**: p<0.01). **(C)** SIRT1 and the PP4 complex cofractionate in fractions around 600kDa (fractions 13-16). Glycerol gradient analysis (12.5%-35%) of nuclear extract from HeLa cells expressing FlagSIRT1 and treated with H_2_O_2_ before harvest (see Methods and Supplementary information). **(D)** HA and FLAG co-immunoprecipitations of HA-PP4 and FLAG-SIRT1, SIRT1 catalytically inactive point mutant H133Y (SIRT1m) or SIRT6 in whole-cell extracts from HeLa cells expressing the indicated constructs treated or not with 2mM H_2_O_2_ for 1h before harvest. **(E)** Time course experiment from 0-24h of SIRT1/PP4 complex assembly upon oxidative stress. PP4c (FLAG) immunoprecipitation of whole-cell extracts from HeLa cells expressing HA-tagged SIRT1 and PP4R2, FLAG-tagged PP4C, and Myc-tagged PP4R3α/β treated with H_2_O_2_ 2mM for 1h and harvested at the indicated times.

SIRT1 enzymatic activity did not regulate this interaction as both SIRT1 WT and catalytically-dead point mutant H363Y (SIRT1m) immunoprecipitated PP4 to a similar extent (Figure 2D). The interaction was SIRT1-specific as we did not detect any interaction between PP4c and SIRT6 under the same conditions (Figure 2D). The SIRT1-PP4 interaction was highly dynamic as it was detected mainly within the 90 min after starting the treatment with H_2_O_2_ (1h incubation+ 30 min after (t_30_)) with a peak at t_0_), and to a lesser degree 6h-12h after the treatment (Figure 2E). Together, this indicated that the SIRT1 and PP4 interaction is enhanced by oxidative stress in SIRT1 activity independent manner.

### SIRT1 inhibits PP4c phosphatase activity

As reported(30), we detected decreased levels of γH2AX in SIRT1-deficient MEFs even 48hr after induction of DNA damage (Figure 3A and Supplementary Figure S2A). Considering that γH2AX is also a relevant target of PP4(12,13), we analyzed the functional implications of SIRT1-PP4 interaction on γH2AX. For that purpose, we purified the PP4 complex (PP4c-R2-R3α-R3β) from HeLa cells and performed *in vitro* phosphatase assays in the absence or presence of SIRT1 using γH2AX as a substrate. Incubation of SIRT1 in presence of NAD^+^ was able to completely repress the phosphatase activity of the complex, suggesting a direct modulation of PP4 activity through deacetylation (Figure 3B). SIRT1 also had a mild inhibitory effect in the absence of NAD^+^ (Figure 3B) and the γH2AX phosphatase activity of PP4 purified from SIRT1-deficient MEFs was significantly higher than when expressed in *Wt* MEFs (Figure 3C). Consistently, the activity of the PP4 complex purified from cells treated with SIRT1 specific inhibitor Ex-527, or with the general Sirtuin inhibitor Nicotinamide (NAM) resulted in a hyperactivation of the phosphatase activity of the complex (Figure 3D). Conversely, the inhibition of PP4 activity with Okadaic acid(37) partially recovered the decreased levels of γH2AX detected in SIRT1-deficient cells compared to Wt cells (Figure 3E) and inhibition of PP4 activity in a SIRT1-deficient background resulted in a milder increase in γH2AX compared to *Wt* (Figure 3E).

**Figure 3:**
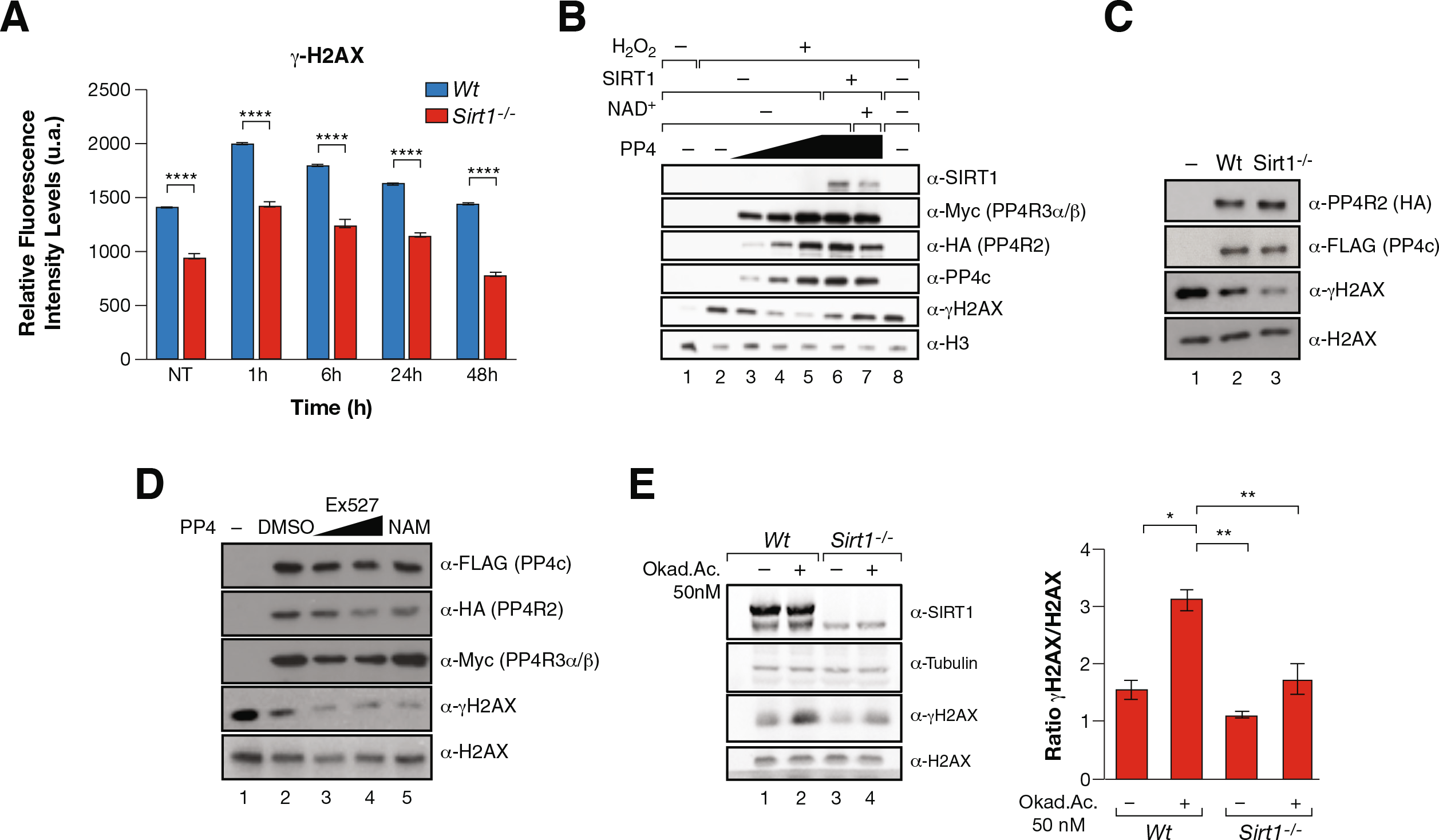
SIRT1 inhibits PP4 phosphatase activity. **(A)** High throughput microscopy (HTM) immunofluorescence analysis of γH2AX in *Wt* and *Sirt1^−/−^* cells treated or not with 7.5Gy IR for 1h and analyzed at the indicated times after irradiation. Data are representative of three independent experiments. Two-tailed t-test (****:p<0.001). At least 300 nuclei were analyzed and the mean with SEM is shown for independent cultures. **(B)** In vitro γH2AX phosphatase activity of a titration of PP4 complexes (1×, 2×, 3×) purified from HeLa cells treated or not with 2mM H_2_O_2_ for 1h and incubated −/+ SIRT1 in presence or absence of NAD^+^. **(C)** *In vitro* phosphatase activity as in (**B**) of PP4 complex purified from Wt and *Sirt1^−/−^* MEFs under oxidative stress 2mM H_2_O_2_ for 1h. **(D)** Assay as in (**B**)-(**C**) Purified PP4 complexes in the presence of EX527(1μM, 10 μM) and 1mM Nicotinamide (NAM). **(E)** Phosphatase assay of whole-cell extracts from Wt and *Sirt1−/−* cells previously treated with Okadaic Acid (OA) (50 nM) or DMSO (control) for 24 hours followed by 1h incubation of 2mM H_2_O_2_ before harvest. Quantification of the ratio of үH2AX/ H2AX in the phosphatase assay is shown.

Altogether, these results suggested that functional interplay between SIRT1 and PP4 regulates γH2AX phosphorylation dynamics.

### SIRT1-dependent deacetylation of K64R in the WH1 domain of PP4R3α and PP4R3β regulates PP4 complex activity

To determine whether the mechanism of PP4 inhibition involved SIRT1-mediated deacetylation, we expressed the PP4 core complex (PP4c-R2) and purified it from *Wt* and *Sirt1^−/−^* MEF cells, resulting in a reconstituted PP4 complex containing all four components PP4 complex (PP4c-R2 and endogenous mouse PP4R3α and 3β). Acetylation analysis of the complex identified two acetylated lysine residues in the PP4R3 subunits in *Sirt1^−/−^* cells compared to *Wt* cells (Figure 4A-C and Supplementary Figures S3A-C); one was common for both PP4R3α and β (K64), and the other ones were specific to each subunit (in K642 for PP4R3α and mouse K777 / human K806 for PP4R3β). K64 is present in the N-terminal WH1 domain, shared by both PP4R3α and β, and is highly conserved in the PP4R3 lineage from *Drosophila* to humans (Figure 4D). The WH1 domain plays a role in protein-protein interactions and has been found in factors involved in cell signaling, cell cycle, cytoskeleton organization, cell motility, gene expression and chromatin organization(38).

**Figure 4:**
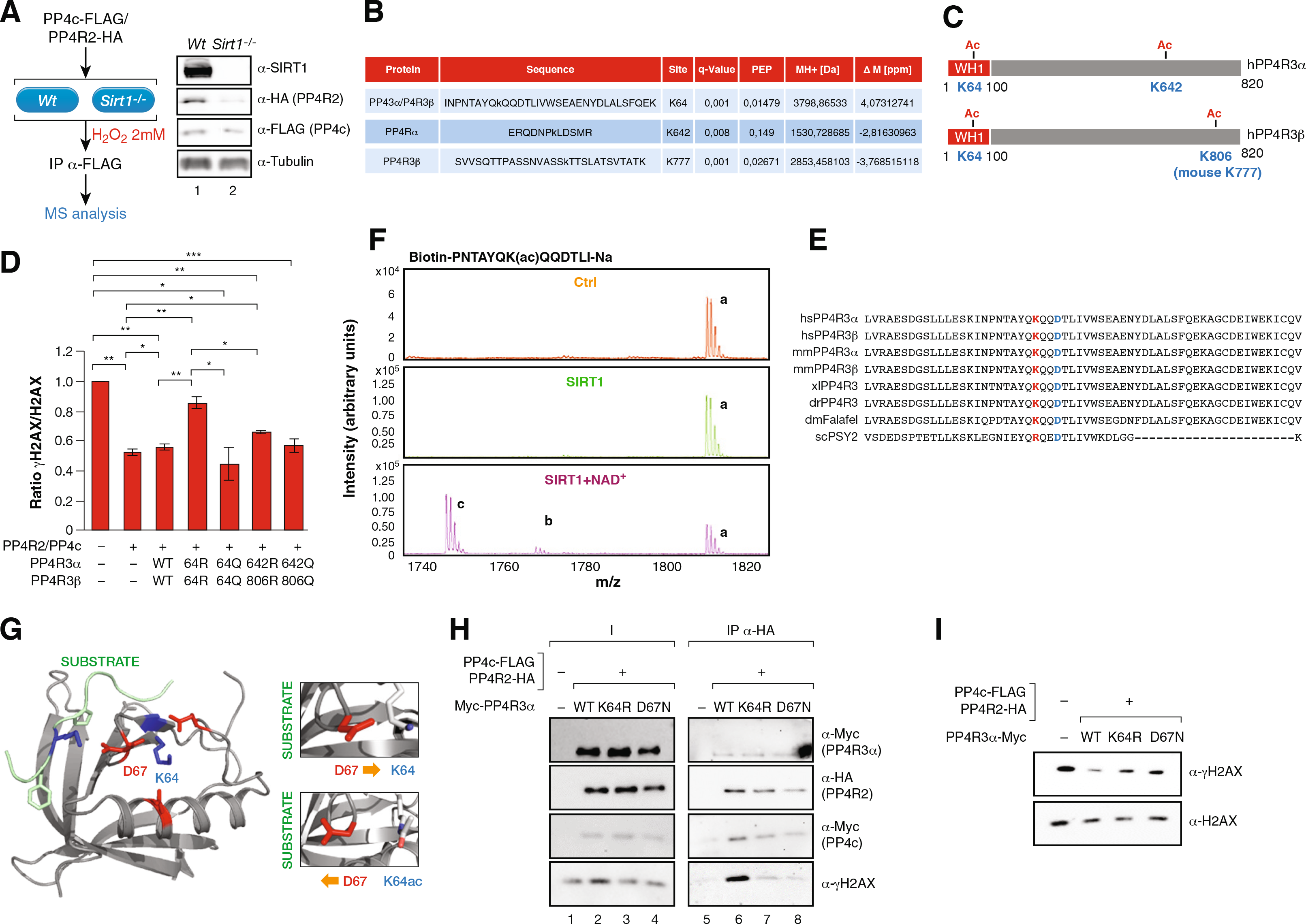
SIRT1 targets K64 in PP4R3α, β to regulate PP4 activity. **(A)**. PP4 complex was purified from MEFs *Wt* and *Sirt1^−/−^* under oxidative stress (2mM H_2_O_2_ 1h) overexpressing PP4 core complex (PP4c and PP4R2) and analyzed by MS to identify acetylated peptides in the PP4 complex upon loss of SIRT1. Left, Schematic representation of the experiment’s pipeline. Right, Levels of the indicated proteins by Western-blot in the MEFs analyzed. **(B)**Summary of the acetylated peptides identified in the MS analysis of A. Differential acetylation was identified in the endogenous PP4R3α and β subunits (see Methods, Supplementary information and Table S2). **(C)**PP4R3α and β protein domains, the acetylation events identified in *Sirt1^−/−^* but not in *Wt* cells, and their position in both subunits. **(D)**K64 and D67 are two conserved residues in WH1 domain of PP4R3α,β from *Drosophila* to humans. WH1 domain primary sequence of the homologs of both PP4R3α,β isoforms in Saccaromyces cerevisiae, Drosophila melanogaster, Xenopus laevis, zebrafish, mouse and humans PP4R3 homologs. **(E)**In vitro үH2AX phosphatase activity assay of human PP4 complex containing the indicated mutations expressed and purified from HeLa cells. A quantification of n=3 experiments (Supplementary Figure S4D) is shown. Two-tailed t-test (*:p<0.05;**:p<0.01;***:p<0.005). **(F)**MALDI-MS analysis of an in vitro K64ac deacetylation reaction by SIRT1. The chromatograms for the control reactions without neither SIRT1 nor NAD^+^ (Ctrl) and with SIRT1 but not NAD+ (SIRT1) are shown together with the complete reaction (SIRT1+NAD^+^). The acetylated peptide was detected either alone or in form of a single sodium adduct. Only the complete reaction rendered deacetylation of both species found for the peptide (b and c) in contrast to the acetylated forms of the peptide (a) (see Methods). **(G)**Structure of human PP4R3α suggests that K64 and D67 are physically very close and have the potential to interact only if K64 is in its unacetylated form (right upper cartoon). Acetylation of K64 could break the interaction between K64 and D67, allowing D67 to turn to the substrate binding region and interact with the substrate through a lysine residue in the FXXP motif (main and right lower cartoon). **(H)**Pull-down experiments with HA resin of previously purified PP4 complexes containing PP4R3α WT/K64R/D67N and core histones. Both the PP4 complexes and the core histones were independently purified from 293Fcells previously treated with 2mM H_2_O_2_. **I**. γH2AX Phosphatase activity of the PP4 complexes in (**H**).

To test whether the deacetylation of any of these residues could explain the SIRT1-dependent inhibition of PP4 complex activity, we mutated these residues to arginine or glutamine to mimic non-acetylated or acetylated lysines, respectively. We purified the PP4 complex containing different combinations of these PP4R3α and β mutants and tested the activity of the complex in an *in vitro* phosphatase assay towards γH2AX. Of the different combinations we tested, we detected a significant inhibition of the phosphatase activity only when the complex contained both PP4R3α and PP4R3β K64R mutants (Figure 4E and Supplementary Figure S3D). No significant effect was detected when the complex contained at least one K64 residue in either PP4R3 α or β (data not shown). The relevance of K64 acetylation in promoting PP4 activity was further supported by the increased activity observed for PP4 complexes containing both K64Q acetyl-mimicking mutants of PP4R3α and R3β (Figure 4E and Supplementary Figure S3D).

To understand SIRT1and K64 acetylation functional relationship, we first confirmed that K64 was a direct target of SIRT1 in an *in vitro* deacetylation assay (Figure 4F). The structure of the human PP4R3α EVH1 domain bound to WAPL through a FXXP motif (6R81, 1.52A) (PMID:31585692) and *Drosophila Melanogaster*’s PP4R3α EVH1 domain bound to CENP-C (4WSF, 1.50A) (PMID 31585692) have been reported. Structural analysis suggests that K64 is not present in the substrate binding site of the domain, but could interact with D67, a conserved residue with the potential to position itself towards the bound substrate, which could result in decreased substrate recognition (Figure 4D and 4G). In contrast, acetylation of K64 would block this interaction, releasing D67 and allowing its repositioning towards the substrate. Supporting a role for D67 in the catalytic activity of the PP4 complex mutation of this residue to Asn (D67N) in PP4R3α, which abrogates the ability to interact with K64 while keeping a similar physicochemical environment, resulted in a decreased ability of this complex to bind to γH2AX-containing histones in pull-down experiments (Figure 4H). A similar effect was observed with the K64R mutant, indicating that the effect of both mutants on PP4 complex are equivalent (Figure 4H). To analyze the effect of these mutations on PP4 catalytic activity we performed γH2AX phosphatase assays with the same complexes tested in the pull-down experiments. Confirming our hypothesis, the results showed that both K64R and D67N mutations significantly decreased the phosphatase activity of the PP4 complex towards γH2AX (Figure 4I).

Altogether, our results demonstrate a functional antagonism between SIRT1 and the PP4 complex upon stress and suggests that SIRT1 inhibits PP4 complex activity towards γH2AX by blocking PP4 substrate recognition through deacetylation of the WH1 domain of regulatory subunits PP4R3α and β.

### SIRT1 antagonizes RPA2 dephosphorylation by PP4

We further analyzed the impact of SIRT1 on the other major PP4 target in the DDR, RPA2. *Sirt1^−/−^* MEFs showed impaired RPA2-foci formation upon H_2_O_2_ or IR treatment without altering overall RPA2 protein levels (Figure 5A-B), suggesting that SIRT1 activity had an impact on RPA2 function. Both *Wt* and *Sirt1^−/−^* cells showed a similar S-phase progression (Supplementary Figure S4A), indicating that RPA2 foci formation was not caused by an altered S-phase. This effect on RPA2 foci formation correlated with a decrease in ph-RPA2 (Figure 5B) and in particular with ph-S33 in *Sirt1^−/−^* cells under IR or H_2_O_2_ treatment (Figure 5C and Supplementary Figure S4B-C, respectively).

**Figure 5:**
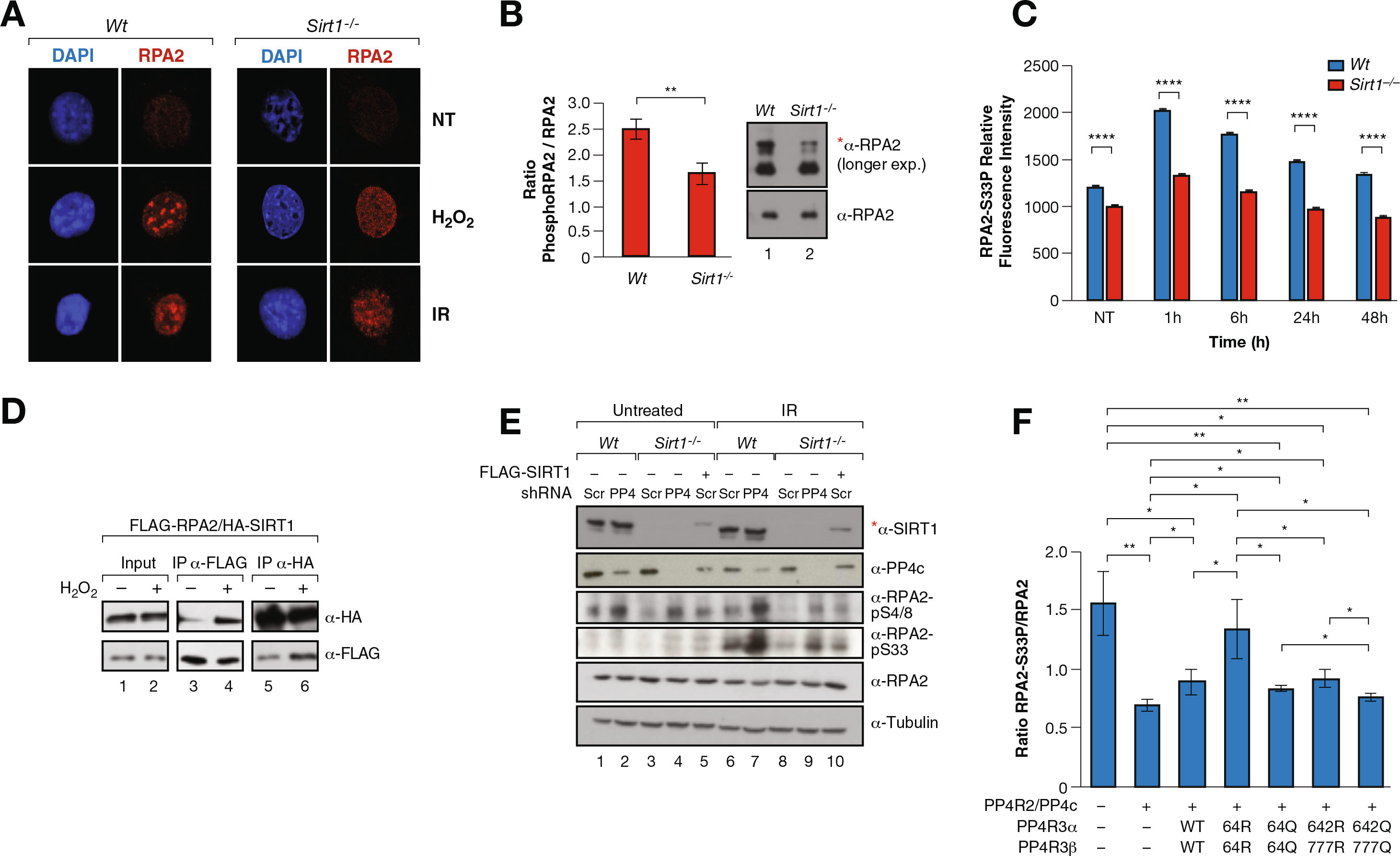
SIRT1 regulates RPA2 phosphorylation through PP4. **(A)** IF analysis of γH2Ax and RPA2 foci formation in Wt or *Sirt1^−/−^* MEFs under or not oxidative stress (2mM H_2_O_2_ for 1h), or ionizing irradiation (IR 30min 7.5Gy). **(B)** Western-blot analysis of the levels of the total pRPA2 in *Wt* and *Sirt1^−/−^* MEFs previously treated with 7.5Gy IR. Quantification of n=3 experiments and representative experiment are shown. Two-tail analysis (**:p<0.01). **(C)** IF Time-course experiment of RPA2-S33P performed as in 3A. A similar experiment is shown by western-blot in Figure S4B. **(D)** Co-IP between FLAG-RPA2 and HA-SIRT1 using FLAG and HA resin under untreated condition or oxidative stress (2mM H_2_O_2_ for 1h) in HeLa cells. **(E)** Levels of the indicated proteins and RPA2 marks upon shRNA-driven downregulation or not of PP4c in *Wt* and *Sirt1^−/−^* MEFs untreated or irradiated with IR (7.5Gy). Lanes 5 and 10, SIRT1 expression was re-introduced in *Sirt1^−/−^* (indicated with *). **(F)** Similar experiment as in 3D with RPA2-S33P levels.

Oxidative stress boosted SIRT1 specific interaction with RPA2 (Figure 5D), and RPA2 co-fractioned with both SIRT1 and PP4 in a high molecular weight complex formed upon stress (Data not shown). In agreement with a key role of PP4 in SIRT1-dependent regulation of ph-RPA2, RPA2 pS33 and pS4/8 hyperphosphorylation induced by PP4 downregulation in *Wt* MEFs was significantly reduced in *Sirt1^−/−^* MEFs (Figure 5E). This decrease was directly dependent on SIRT1 protein as re-expression of SIRT1 partially recovered the PTM levels (Figure 5E). Similar to the case of γH2AX, the K64R mutation in both PP4R3α/β resulted in increased levels of RPA2 phosphorylation, while K64Q had the converse effect (Figure 5F and Supplementary Figure S4D), supporting that regulation ofpRPA2 by SIRT1 was associated with the same mechanism of PP4 regulation through PP4R3α/β. Taken together, these results confirmed a direct role of SIRT1 in the dynamics of RPA2 phosphorylation through the regulation of PP4 complex activity.

### Interplay between SIRT1 and PP4 regulates HR and genome stability

To determine the functional impact of the interplay between SIRT1 and PP4 we analyzed cell cycle progression and DNA repair kinetics. While downregulation of PP4c in *Wt* MEFs resulted in an increase of cells in G_1_ phase and a decrease in S-phase, it did not alter significantly the Sirt*^1−/−^* MEFs cell cycle (Figure 6A and Supplementary Figure S5A). A similar result was observed upon downregulation of PP4c and/or SIRT1 in U2OS cells using siRNAs (Supplementary Figure S5B). The increase in G_1_ induced by PP4c downregulation corresponded to mild but significant decrease in cell proliferation that was rescued by SIRT1 depletion (Supplementary Figure S5C). Notably, cell viability was no affected under any condition (Supplementary Figure S5D).

**Figure 6:**
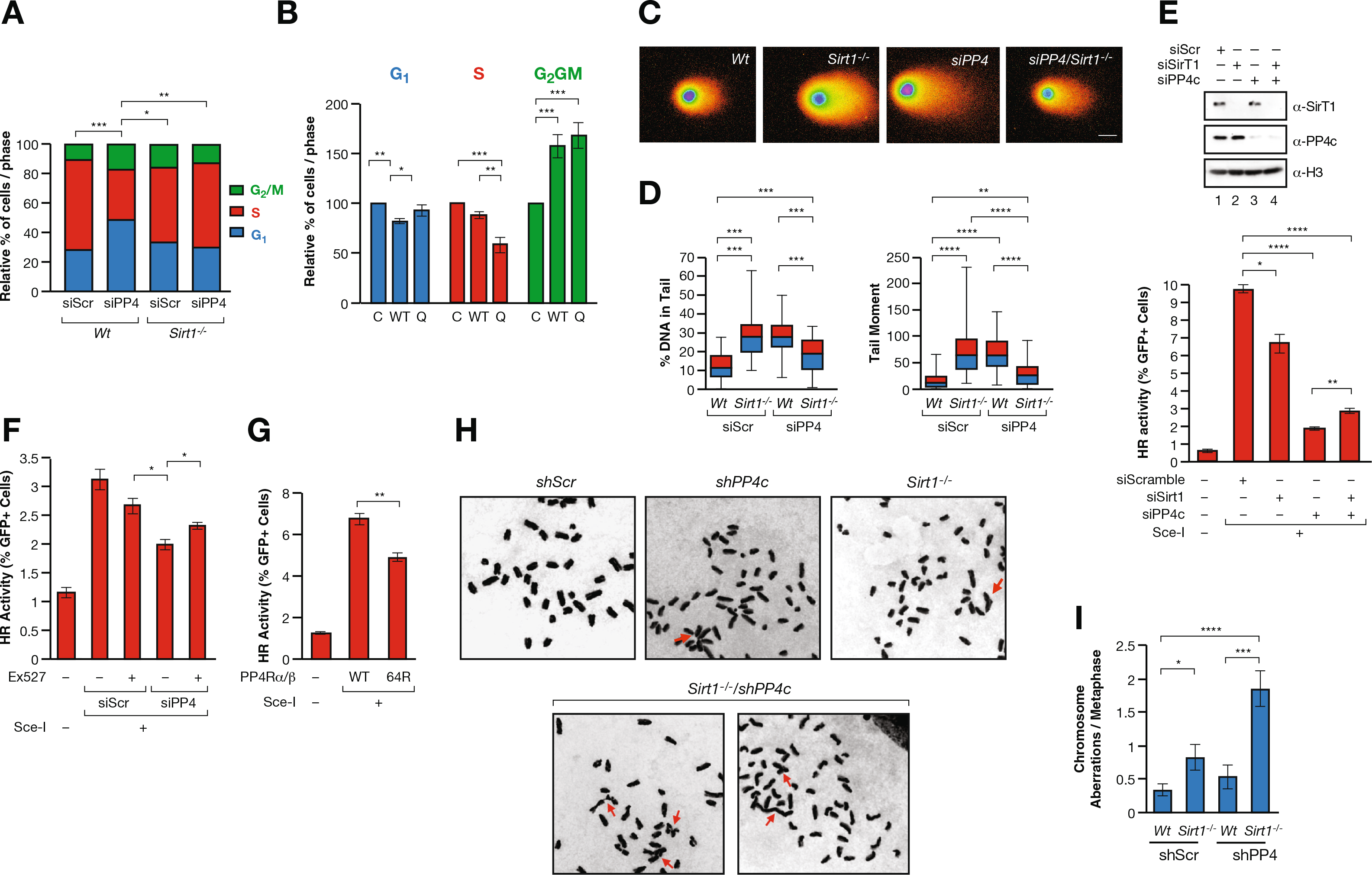
The interplay between SIRT1 and PP4 regulates DNA repair and genome stability. **(A)** Quantification of Propidium iodine (PI)/EdU based cell cycle analysis (n=3) of MEFs *Wt* or *Sirt1^−/−^* transduced with shRNAs scramble or targeting PP4 (shPP4). **(B)** Cell cycle analysis as in (**A**) of U2OS cells transfected with an empty (C), wild-type PP4R3α,β (WT) or mutant PP4R3α,β K64Q (Q) expression vectors **(C)-(D)** Neutral comet assay of U2OS cells *Wt* or *Sirt1^−/−^* expressing siRNA scramble or against PP4 treated with of 1μM Camptothecin for 90 min. A representative full-spectrum image of a cell of each condition **(C)** and quantification of n=3 experiments **(D)** are shown. The % DNA in tail and tail moment are quantified. A similar analysis in absence of Camptothecin is included in Supplementary Figure S5E. Two-tailed T-test assay was performed (**:p<0.01; ***:p<0.005; ****:p<0.001). **(E)** *in vivo* HR reporter system in U2OS cells upon siRNA-driven downregulation of PP4c and/or SIRT1. Western-blot of SIRT1 and PP4c levels in each condition (top) and. quantification of n=3 experiments(bottom) are shown. Two-tailed t-test analysis was performed(*:p<0.05; ***:p<0.005; ****:p<0.001). **(F)-(G)** HR assays as in (**E**) upon treatment with either DMSO (control) or 10μM of SIRT1 inhibitor Ex-527 (**F**) or upon overexpression of both PP4R3α and β WT or K64R (**G**). **(H)-(I)** Chromosome aberration analysis of the same cells as in (**A**) treated with 250nM of Mitomycin C for 24h. Representative images of each condition (**H**) and a quantification of n=3 experiments (**I**) are shown. Two-tailed t-test analysis was performed(*:p<0.05; ***:p<0.005; ****:p<0.001).

We next tested the effects of PP4R3α/β WT or K64Q overexpression on cell cycle progression. We observed that while both WT and K64Q induced an increase in G_2_/M cells, we only observed a clear and significant differential effect between PP4R3α/β WT and K64Q in S-phase (Figure 6B), suggesting that the SIRT1/PP4 interplay is primarily relevant during S-phase progression. We next analyzed DSB-associated DNA damage by neutral comet assay upon induction of S-phase specific replicative damage with the Topoisomerase-1 inhibitor campothecin (CPT). The results confirmed that while downregulation of either SIRT1 or PP4 induced increased levels of DSBs, downregulation of both resulted in a significant compensatory effect (Figure 6C-D and Supplementary Figure S5E).

As both PP4 and SIRT1 have been implicated in HR, we measured the impact of their depletion on HR DNA repair using a well-established *in vivo* HR reporter in U2OS cells, DR-HR(39). Following DNA damage induction by the nuclease Sce-I, efficient HR results in a reconstitution of intact GFP that can be monitored by flow cytometry (Supplementary Figure S5F). We observed that while siRNA-driven downregulation of SIRT1 resulted in a roughly 33% decrease in the HR capacity of these cells, downregulation of PP4c had a drastic effect on HR efficiency, which dropped to 20% of the observed with the siScrcontrol (Figure 6E and Supplementary Figure S5G). Supporting the functional antagonism between SIRT1 and PP4, downregulation of both PP4c and SIRT1 resulted in a significant partial rescue of the HR efficiency relative to the single downregulation of PP4c. The mild recovery of the HR efficiency upon loss of both PP4c and SIRT1 may indicate a dominant role of PP4 in the process that includes functions that are not regulated by SIRT1 activity. Chemical inhibition of SIRT1 with EX-527 caused a similar reduction of the effect of PP4c downregulation on HR (Figure 6F). We confirmed that K64 acetylation regulation is involved in HR regulation, as overexpression of the PP4R3α/β K64R mutants in these cells significantly decreased the HR activity in around 35% compared to WT proteins, strongly supporting a role for PP4R3α/β deacetylation in the regulation of HR (Figure 6G).

To study the functional consequences of this interplay we analyzed the levels of genomic instability following their downregulation and treatment with mitomycin C (MMC), a DNA crosslinker that promotes general double strand breaks throughout the cell cycle. We then measured the number of chromosomal aberrations in metaphase spreads. Surprisingly, while loss of SIRT1 or shRNA-driven downregulation of PP4c resulted in an increased number of chromosomal aberrations, the combined effect of both resulted in a significant higher number of aberrations (Figure 6H-I). These results suggested that the interplay between both factors may go beyond HR in S-phase. A possible explanation is that loss of both factors not only induce an increase in HR efficiency, but also of other pathways with lower accuracy in the repair, such NHEJ. Of note, we discarded that our observations are related to cell death, as we did not detect an increase in apoptotic cells in any of these treatments (Supplementary Figure S5H). Nevertheless, the fact that SIRT1 loss in cells increases their sensitivity to PP4c downregulation strongly supports a functional antagonism between both factors in genome stability.

### SIRT1 and PP4 levels inversely correlate in vivo in breast cancer

Given the direct impact of DNA damage and genome stability in human health, we hypothesized that SIRT1 and PP4 may perform opposing roles in human pathologies associated with genome instability, such as cancer. To study the impact of this interplay *in vivo*, we analyzed the degree of correlation between SIRT1 and PP4c levels in a cohort of 94 breast cancer samples from the National Cancer Institute’s Clinical Proteomic Tumor Analysis Consortium (CPTAC), where a complete proteomic/phosphoproteomic analysis had been performed (Figure 7A). We selected breast cancer in our study as it has a well-established link to genome stability accumulation and DNA repair defects (40). As expected, we observed a general inverse relationship between SIRT1 and PP4c protein abundance (Figure 7B) while SIRT6 showed a direct correlation with PP4c in these tumors. Consistently, the levels of pRPA2(p238) and another PP4 target, pTRIM28(pS473), directly and inversely correlated with SIRT1 and PP4, respectively (Figure 7C and Supplementary Figure S6A). In contrast, neither SIRT1 nor PP4c correlated with other phosphorylation events unrelated to PP4, such as pATM (Supplementary Figure S6B). We next analyzed whether this correlation was valid in the subset of samples with mutations or deletions in genes associated with Homology directed repair (HDR) or PARP inhibitor sensitivity (Figure 7D). Although this filter considerably reduced the number of samples, SIRT1 and PP4C levels showed an anti-correlation at both the protein (Figure 7E) and RNA level (Supplementary Figure S6C). Altogether, this evidence further supports an *in vivo* specific antagonism between SIRT1 and PP4 in tumorigenesis, reflecting the functional relevance of the interplay between both factors in the maintenance of genome stability.

**Figure 7:**
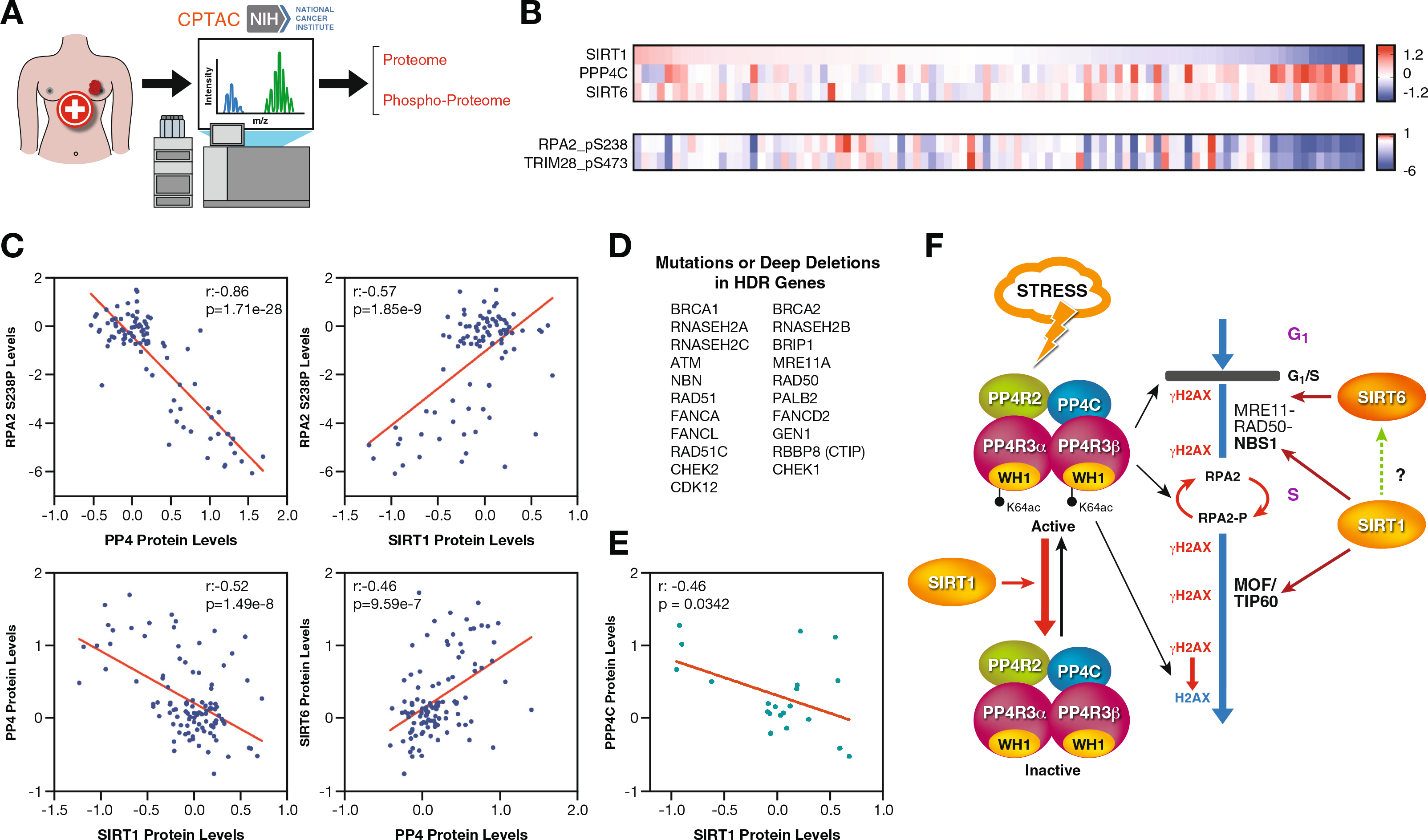
SIRT1 and PP4 inversely correlate in cancer. **(A)** Proteomics and phosphoproteomics data from tumor samples was obtained from the National Cancer Institute’s Clinical Proteomic Tumor Analysis Consortium (CPTAC) analysis of tissues from the Breast Invasive Carcinoma (TCGA) dataset. **(B)** Heatmap of SIRT1, SIRT6, PPP4C protein abundance and pRPA2 (p238) and pTRIM28 (pS473) levels in samples from the study described in (**A**). Each column corresponds to one tumor sample. **(C)** Correlations between the indicated proteins and pRPA2 in each tumor sample were analyzed. Pearson correlation coefficient (r) and p-value (p) for each analysis are shown. **(D)** List of HDR-associated genes used in the analysis of (**E**). **(E)** Samples of the Breast cancer cohort containing mutations or deep deletions in the genes in (**D**) were selected and analyzed as in (**C**). **(F)** Proposed model of the antagonistic interplay between SIRT1 and PP4 in HR repair.

## DISCUSSION

Despite the well-established link of Sirtuins with the genome stability, we still have a poorly integrated view regarding how these enzymes regulate the DDR. One of the most intriguing observations, the decrease in γH2AX observed in *Sirt1^−/−^* mice(30) has remained unaddressed. Our work not only provides a mechanism to explain this early observation, but also suggests a more global perspective about how SIRT1 regulates DDR signaling. We hypothesize that in addition to a specific modulation of HR DNA repair signaling at different stages (NSB1, MOF, TIP60)(41,42), SIRT1 regulates the progression of the whole signaling pathway through a direct control of the PP4 complex function (Figure 7E). In the case of γH2AX, we propose that while stress is active, SIRT1 binding to PP4 ensures that the DDR signaling remains on until completion of the HR repair and the stress signal disappears. This effect is likely unrelated to the described deacetylation of NBS1 by SIRT1 (42), as NBS1 do not seem to have an impact in the deposition of the mark(7,43).

In contrast to PP2A, loss of PP4 induces increased levels of γH2AX in absence of external insults, suggesting that PP4 has a key role in basal repair, probably associated with replicative stress. This is further supported by the described antagonism between ATR and PP4(12,44). Interestingly, the link between SIRT1 and replicative stress, and the observed decrease of γH2AX in *Sirt1^−/−^* MEFs also in the absence of an external stimuli, further supports this link. The SIRT1-PP4 association under oxidative stress suggests that they play a prominent role in the regulation of genome stability under these conditions. In this sense, the reported phosphorylation of SIRT1 by JNK1 under oxidative stress may be an important regulator of this mechanism as it promotes its activity and its nuclear localization (62).

Another interesting observation is the striking opposed correlations between SIRT1 or SIRT6 and PP4 and its targets in breast cancer samples(15,45,46) (Figure 7A-E and Supplementary Figure S6). This evidence supports a differential role for both SIRT1 and SIRT6 in DSB repair and may suggest an unexpected functional interplay between SIRT6 and PP4. SIRT1 has been recently reported to interact and deacetylate SIRT6 in K33ac regulating its binding to γH2AX and chromatin (63). However, the fact that SIRT6 was not detected as a component of the SIRT1-PP4 complex (Figure 1), or SIRT6 and PP4 do not seem to interact (Figure 2D) suggests that this correlation is indirect and probably reflects a coordinated regulation of both factors in DDR process. Further studies should define this functional relationship and determine whether SIRT6 has any impact on SIRT1 control of PP4.

Despite the relevance of the interplay PP4-SIRT1, our data also suggest that SIRT1 regulates γH2AX through a PP4-independent mechanism, since increased γH2AX observed upon PP4 downregulation or chemical enzymatic inhibition with Okadaic acid, is significantly decreased in a *Sirt1^−/−^* background (Figure 3E). This is probably caused by a direct or indirect regulation of ATR or ATM activation by SIRT1, which is not well understood. For instance, while SIRT1 recruitment to DNA damage regions depends on ATM and a direct role of SIRT1 in ATM activation has been shown in post-mitotic neurons, ATM/ATR phosphorylation of the SIRT1 inhibitor DBC1 promotes the inactivation of SIRT1 upon DNA damage(33). Furthermore, SIRT1 deacetylates TopBP1, an ATR activator under replicative stress(47), an event that inhibits ToBP1 binding to Rad9 and its subsequent activation of the ATR-Chk1 pathway(48). Further studies should clarify the mechanism involved in this observation.

Considering the observed compensatory effect between SIRT1 and PP4 downregulation at the level of comet assay and HR activity, the observed increase in the frequency of aberrations in metaphase chromosomes upon DSB induction is intriguing. This increase may be explained by the fact that SIRT1 and PP4 also induce NHEJ, the less accurate major DSB repair pathway(49,50). Interestingly, although the main role of both PP4 and SIRT1 in DSB repair is through HR, they have been shown to target KAP1 (SIRT1 and PP4) and 53BP1 (PP4) to promote NHEJ activity(49,51,52). However, other evidence suggests that PP4 has an inhibitory effect on the pathway (50) which would fit with NHEJ induction upon SIRT1/PP4 downregulation. Moreover, these aberrations were detected upon induction of DSB throughout the cell cycle with mitomycin-C, which may suggest that part of these aberrations may be due to non S-phase damage, maybe during G_1_ or G_2_/M. Thus, some studies suggest that PP4 plays a role in the deposition of 53BP1 pathway during mitosis for NHEJ repair in G1, which may support this possibility(52).

Interestingly, H4K16ac deacetylation by SIRT1 was also decreased upon incubation with the PP4 complex (Supplementary Figure S2B), which suggests that PP4 modulates SIRT1 activity. We speculate that it may involve two different mechanisms: First, the dephosphorylation of SIRT1, may play a key role, as phosphorylation by stress-dependent kinases like JNK1, ATM, AMPK or MAPK was shown to be essential for SIRT1 activation upon stress (53,54). The reports that phosphorylated SIRT1 localizes to replication origins to prevent excess origin firing(55) while PP4 dephosphorylation of yeast initiation factors promotes origin firing, may suggest that the SIRT1/PP4 interplay may also regulate DNA replication. Whether this is also valid in human origin activation is currently unknown(56). We hypothesize that the resumption of cell cycle progression after DNA repair, may involve SIRT1 inactivation by PP4. Second, PP4 may also regulate SIRT1 activity through another factors. Thus, some studies have identified the SIRT1 inhibitor DBC1 as a target of PP4(33) and this dephosphorylation may induce binding of DBC1 to SIRT1, blocking its activity.

The functional implications of the SIRT1/PP4 interplay in regulation of RPA2 phosphorylation are less clear. While phospho-RPA2 is involved in specific binding to DNA repair factors, its hyper-phosphorylation inhibits RPA complex binding to DNA, delays the formation of RAD51 foci, and blocks DNA replication(25). Interestingly, RPA2 dephosphorylation by PP4 is required to complete DNA repair and to resume replication after DNA damage is completed(22). In this sense, like in the case of γH2AX, the increase of RPA2 phosphorylation by SIRT1-dependent inhibition of PP4 could ensure the activation of the pathway until DNA repair is completed. However, the established role of PP4 in RPA2 regulation and our observation that SIRT1 deficiency also results in a decrease in RPA2 foci formation (Figure 5c), suggests that the dynamic regulation between both factors is key to promote efficient repair of damaged DNA regions.

Another interesting discovery from our work is the identification of PP4R3α and β as the mediators of SIRT1’s regulation of PP4 activity. The identification of K64 in the WH1/EVH1 domain of PP4R3α,β as SIRT1 target is particularly interesting giving the key role of this domain in the binding to the PP4 substrates(38). The functional crosstalk between SIRT1 and PP4 is likely to involve many other physiological processes, although is not clear whether this is based on a functional antagonism between both factors like in the case of DDR. An obvious example is oncogenesis as PP4 has been described to be overexpressed in several types of cancer, among which is breast cancer(15,16,57,58) while SIRT1 plays a more complex role with tumor suppressor or pro-oncogenic roles depending on the context. since our evidence suggests that both factors could play antagonistic roles in breast cancer, determining the implication of PP4 in any of SIRT1 specific roles in cancer or vice versa could further support the functional implications of this interplay. Further studies should validate these observations.

Altogether, our work not only suggests a novel regulatory mechanism by SIRT1, but also proposes a new perspective about the Sirtuin-dependent control of DDR where they exert a global control of the process by modulating the cascade of phosphorylation events involved. This mechanism may have further biological implications and reflect a more complex interplay between SIRT1 and the PP4 complex. Future studies should determine the extent of this functional relationship and its relevance in SIRT1-dependent functions.

## Supporting information

Supplemental_Information_Rasti_et_al

## DATA AVAILABILITY

The mass spectrometry data generated in this work is included in the main figures and supplementary tables S1 and S2. Mass spectrometry analysis of PP4 complex acetylation has been deposited to the ProteomeXchange Consortium via the PRIDE partner repository database (http://www.ebi.ac.uk/pride) with the data set identifier PXD036905.

## SUPPLEMENTARY DATA

Supplementary Data are available at NAR online

## ACKNOWLEDGEMENTS

We are grateful to Dr. Anne-Claude Gingras (LTRI, Toronto) for providing PP4c expression vector and Dr Ranjit S Bindra (Yale Univ, New Haven) and Dr Simon N. Powell (MSKCC, NYC) for providing the DR-HR U2OS cell line. Thanks to the members of the Vaquero laboratory for stimulating discussions.

## FUNDING

This work was supported by the Spanish Ministry of Economy and Competitiveness (MINECO) (SAF2014-55964R, SAF2017-88975R, PID2020-117284RB-I00) (AV), PGC2018-095616-B-100/GINDATA (THS) and cofounded by FEDER funds/European Regional Development Fund (ERDF)-A Way to Build Europe; The European Commission’s Horizon 2020 research and innovation programme - Marie Słodowska-Curie grant # 895979 (BNV); The Catalan Government Agency AGAUR (2014-SGR-400, 2017-SGR-148) (AV); Centres of Excellence Severo Ochoa award (THS); and the La MaratÓ de TV3 Foundation (AV). We also thank the CERCA Programme/ Generalitat de Catalunya for institutional support. The work was also supported by a Beatriu de Pinos fellowship AGAUR (BNV); MINECO FPI BES-2015-071251(MEA); FI-AGAUR 2021-FIB1-00228 (AG-G); and “la Caixa” Foundation LCF/PR/GN14/10270002 (SSB).

## CONFLICT OF INTEREST STATEMENT

The authors declare that they have no competing interests.

## Notes

### Competing Interest Statement

The authors have declared no competing interest.

### Summary of Updates

We have acknowledged a project that was not mentioned by mistake.

