## Supplemental_Information_Rasti_et_al for "SIRT1 regulates DNA damage signaling through the PP4 phosphatase complex"

SUPPLEMENTARY INFORMATION

SUPPLEMENTARY FIGURES

A

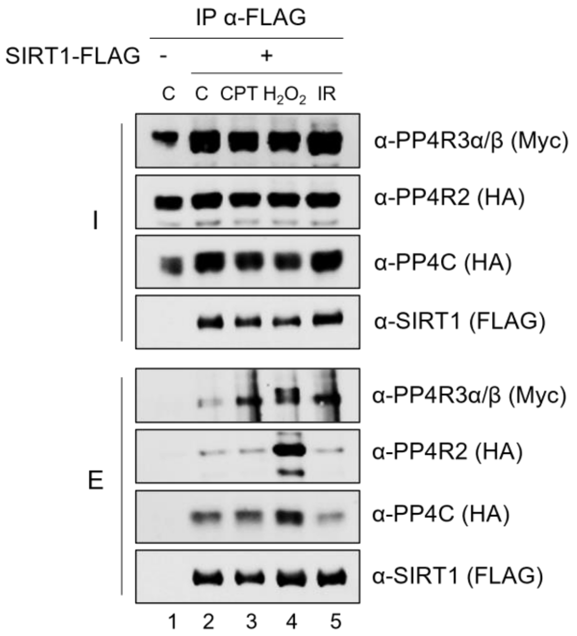


B

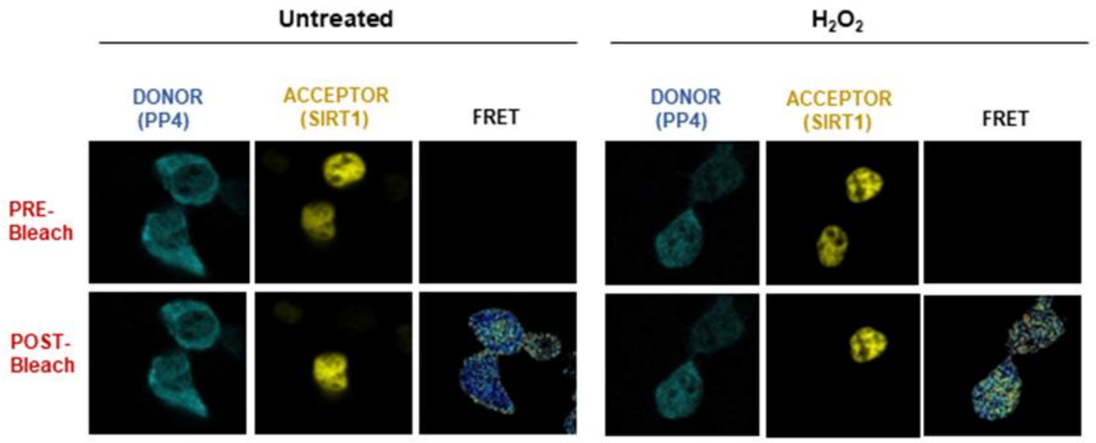


**Supplementary Figure S1** (**A**)**.** Interaction as in Figure 2A of the interaction between SIRT1 and the PP4 complex upon different conditions of stress including 2 mM H2O2 for 1h, 1 uM CPT for 1h and 7.5 Gy IR. (**B**) Representative images of the FRET analysis shown in Figure 2B.

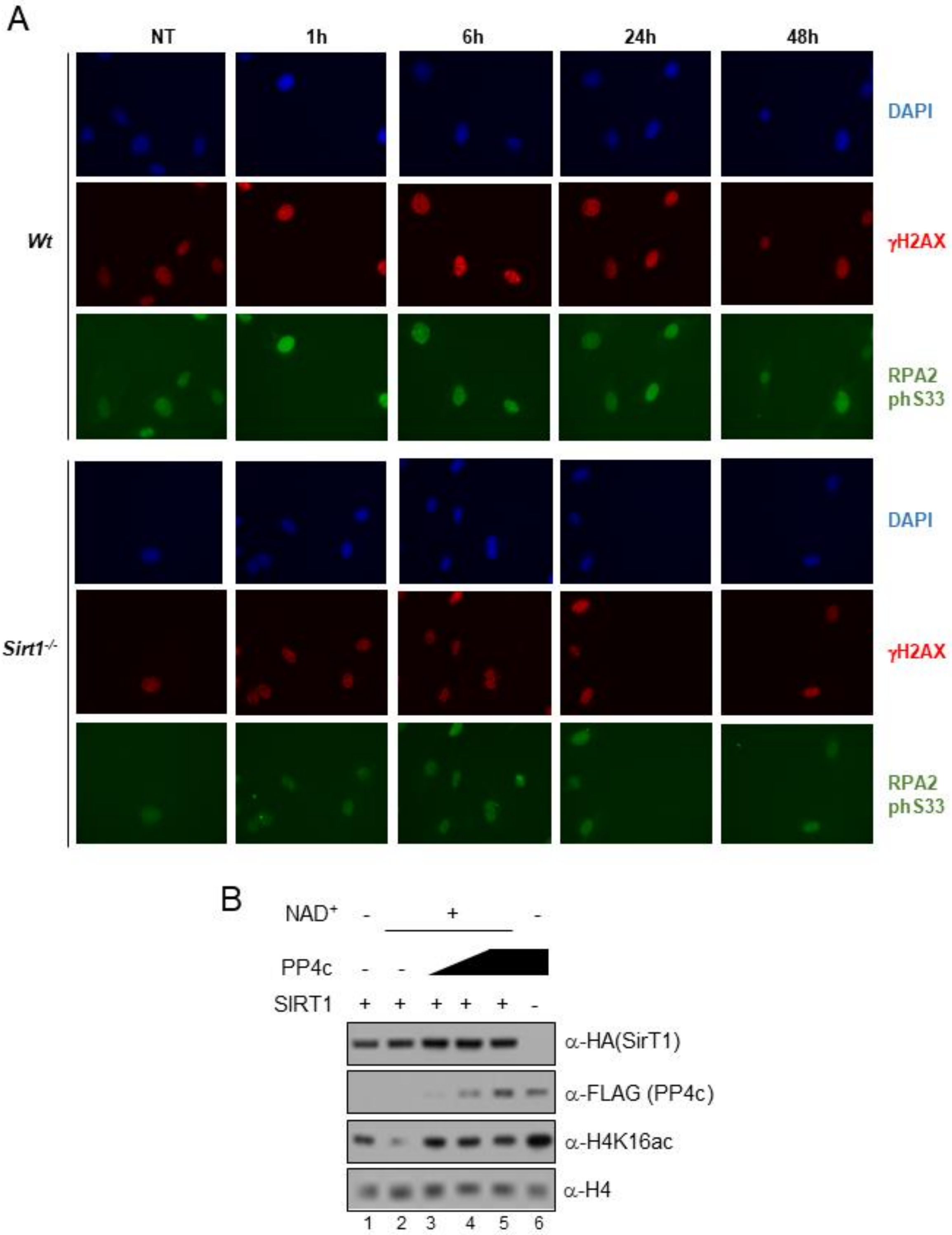


**Supplementary Figure S2**. (**A**) Time course immunofluorescence analysis by High throughput microscopy (HTM) of *Wt* and *Sirt1*$^{-/-}$ MEFs after irradiation or not with 7.5Gy IR for 1h. DAPI, γH2Ax and phospho-RPA2 (S33) were analyzed. (**B**) SIRT1 *in vitro* deacetylase assay. The enzymatic deacetylase activity of SIRT1 towards H4K16ac was tested in presence or absence of increasing amounts of purified PP4 complex as in Figure 3B. Deacetylase reactions were followed by immunoblotting and probed with the indicated antibodies.

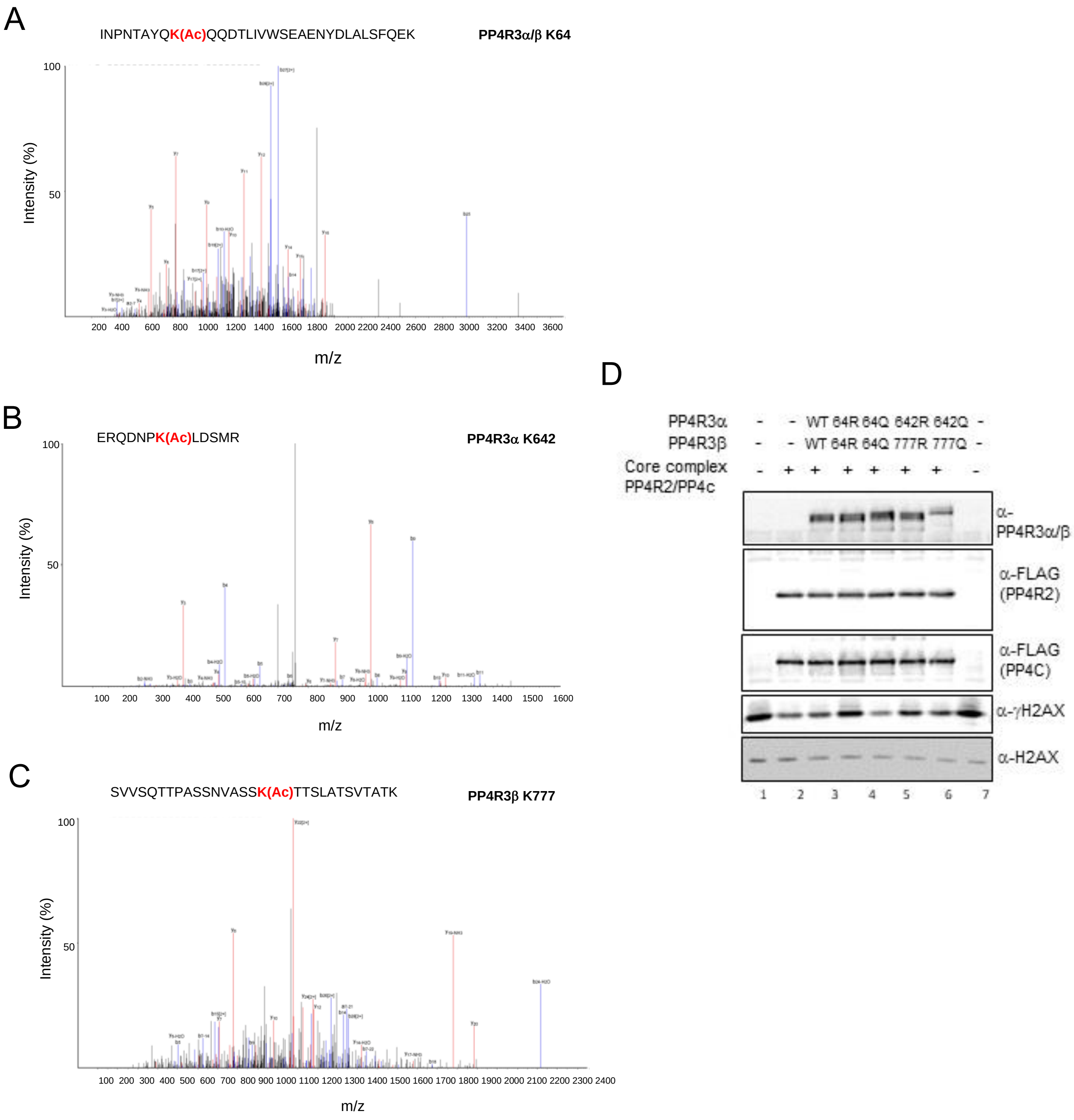


**Supplementary Figure S3**. (**A**)-(**C**) Collision-induced dissociation (CID) spectrum of acetylated peptides identified in the PP4 complex (K64 in α/β, K642 in PP4R3α and K806 in PP4R3β) in the analysis described in Figure 4A-C. (**D**) Representative experiment on n=3 represented in Figure 4E.

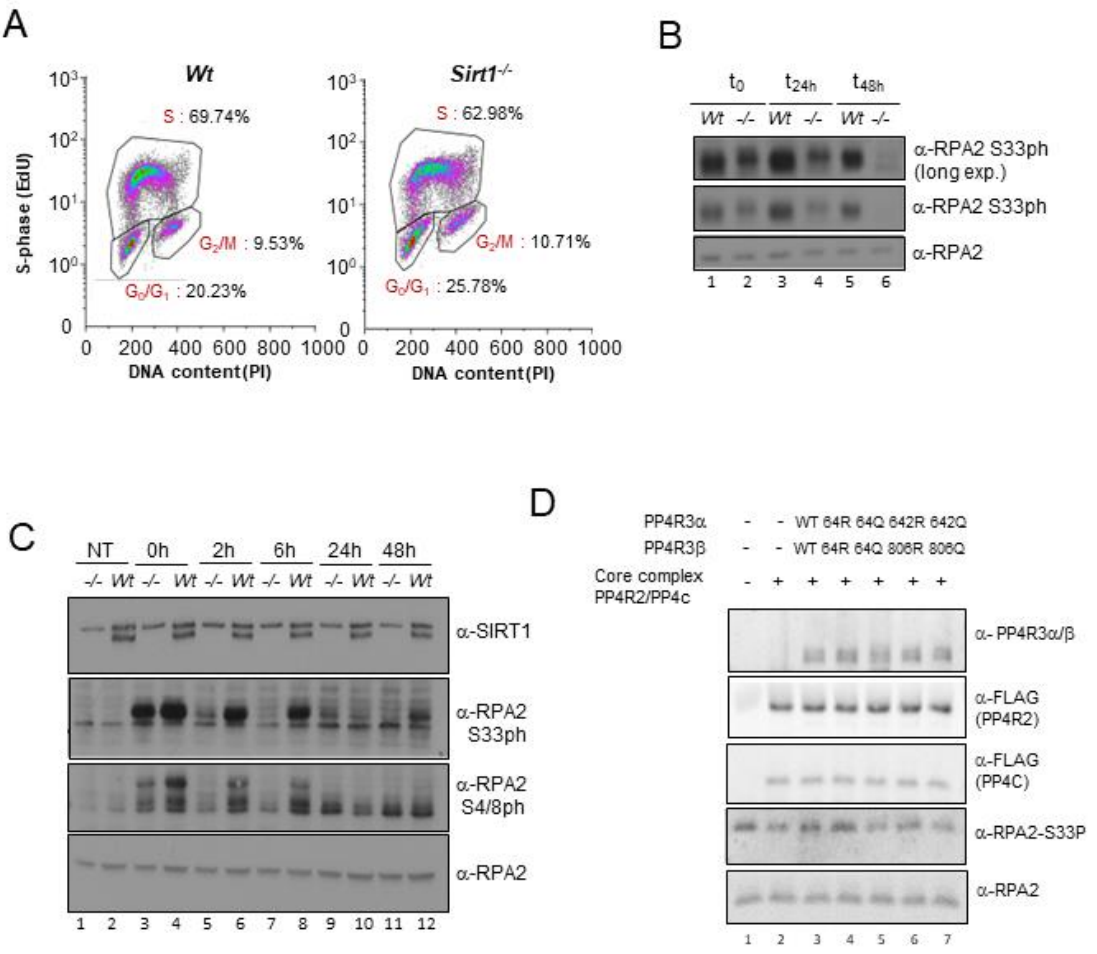


**Supplementary Figure S4**. (**A**) Cell cycle analysis of the MEFs *Wt* and *Sirt1*$^{-/-}$ shown in Figure 5A with double staining of propidium Iodide (PI) and EdU (S-phase specific staining). The results showed a mild decrease in the cells in S-phase. (**B**) Time course of experiment similar to Figure 5C. *Wt* and *Sirt1*$^{-/-}$ MEFs were treated by IR (7.5 Gy) and recovered at 0, 24 and 48 hours after treatment. The total levels of RPA2 and Phospho-RPA2 (S33) were tested by indicated antibodies. (**C**) Similar experiment as in Figure 5C and in (B) *Wt* and *Sirt1*$^{-/-}$ MEFs but under oxidative stress (2.5mM $H_2O_2$) instead of IR. RPA2, phospho-RPA2 (S33 and S4/8) were detected by indicated antibodies. (**D**) Representative experiment on n=3 represented in Figure 5F.

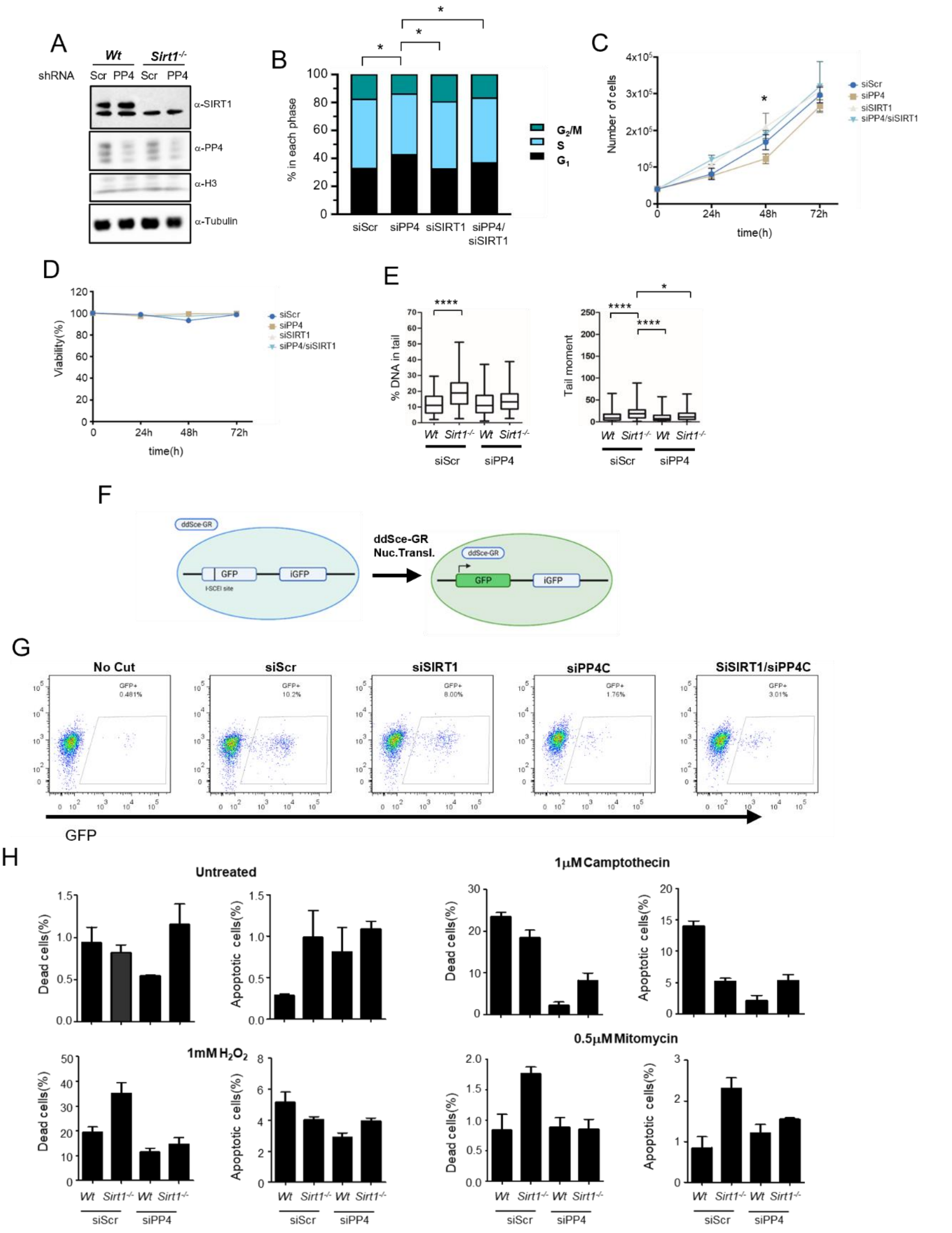

**Supplementary Figure S5**. (**A**) Western-blot analysis of the cells tested in Figure 6A to monitor the levels of SIRT1 and PP4c. (**B**) SIRT1 and/or PP4C expression in U2OS cells were targeted using siRNA and cell cycle distribution was analyzed 48h after siRNA transfection. Two-tail t-test (*:p<0.05). (**C**)-(**D**) U2OS were treated as in (B) and the total number of cells (C) or the proportion dead/alive cells were counted at 24h, 48h and 72h after siRNA transfection. (**E**) Neutral comet assay performed as in Figure 6C-D but in this case in absence of stress. (**F**) Schematic representation of the *in vivo* HR reporter system performed in U2OS cells in Figure 6E-G. (**G**) Representative experiment of n=3 HR assays quantified in Figure 6E. the proportion of detected GFP positive cells are indicated. (**H**) Percentage of cell death and apoptosis upon the indicated treatments in U2OS cells *Wt* and *Sirt1*$^{-/-}$ downregulated or not in PP4c by siRNAs.

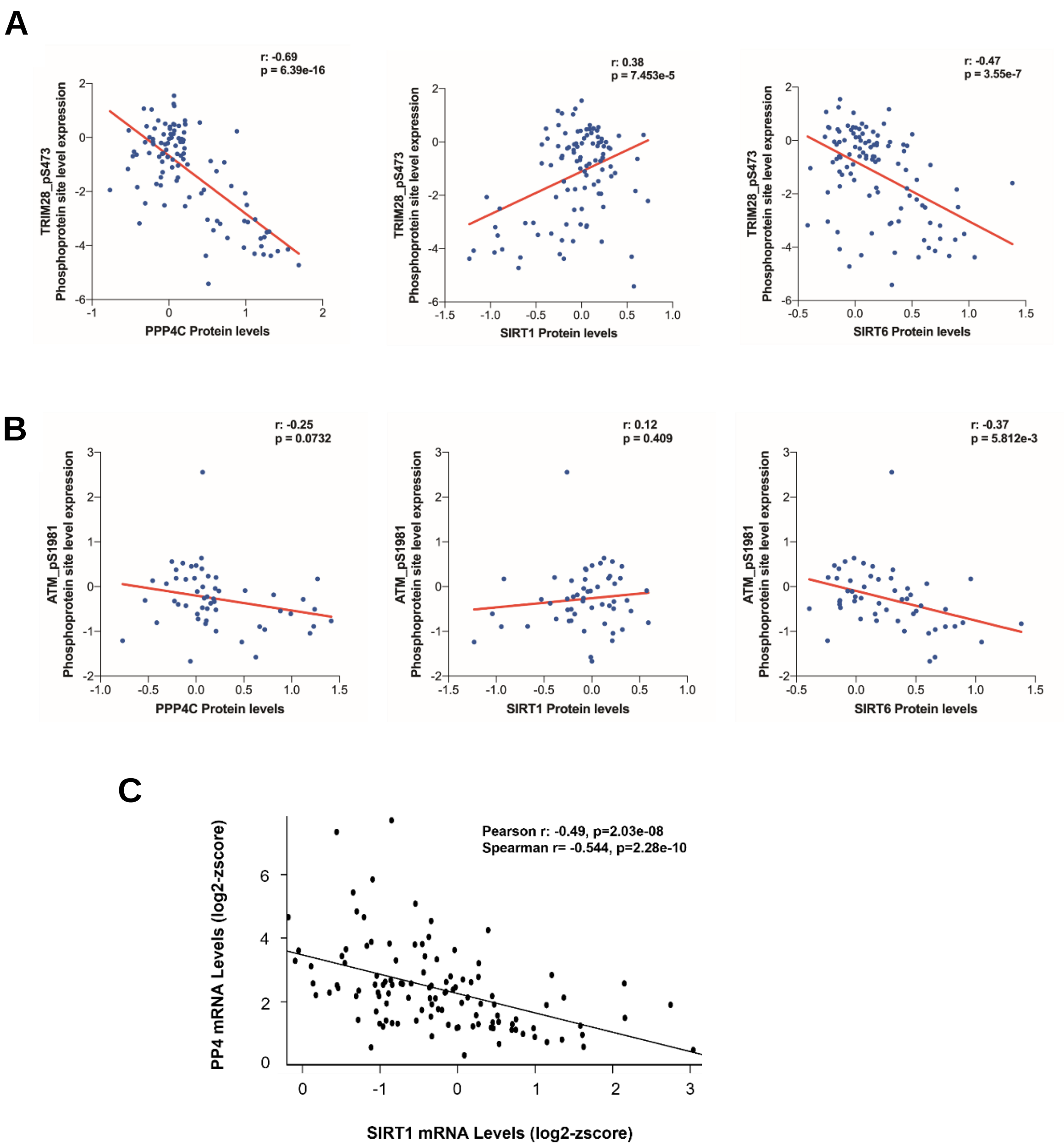


**Supplementary Figure S6**. (**A**) Correlations as in Figure. 7C between pTRIM28 (p473) and either PP4C, SIRT1 or SIRT6 in each tumour sample. Pearson correlation coefficient (r) and p-value (p) for each analysis are shown. (**B**) Similar analysis as in (A) but with pATM (S1981) instead of pTRIM28. (**C**) Gene expression analysis was assessed in the HDR-deficient subset (as defined in Figure 6D) of the TCGA Breast cancer cohort. The correlation between SIRT1 and PP4C is shown.

### SUPPLEMENTARY MATERIALS AND METHODS

#### Polymerase chain reaction (PCR)

PCR reactions were performed using 5-10ng of template DNA in 25µl of GoTaq mix, 0.3µM of each primer and filled with H2O (Milli-Q) up to 50µl. DNA was amplified with 35 cycles of denaturation at 95°C for 45sec, annealing at 57, 59 or 61°C for 60sec, extension at 73°C for 60sec per kilobase. The final extension was performed at 73°C for 5min.

#### Cloning

Vectors and DNA inserts were digested with appropriate restriction enzymes and separated by horizontal electrophoresis. DNA bands were excised from gel and purified with NucleoSpin® Gel and PCR Clean-up kit (Macherey-Nagel, Düren, Germany). For cloning of PCR products, restriction sites were introduced at the ends of primers. The primers used in PCR reactions were synthesized by SIGMA. Fragments were quantified using NanoDrop instrument (Wilmington, USA), subjected to appropriate restriction enzyme digestion and ligated using 1U of T4 DNA ligase in 13µl reaction using 100µg of vector and a corresponding stoichiometric amount of insert at a vector: insert ratio of 1:0, 1:3 and 1:5. Ligations were incubated O/N at 16°C and used to transform 100µl of competent DH5α cells. Some transformed clones were selected for plasmid purification in order to sequence the DNA insert and verify the nucleotide sequence. A single positive colony was grown in LB medium at 37°C with shaking and used for maxiprep. The pcDNA3-FLAG-PP4C was a gift from Dr.Gingras( Gingras et al., 2005).

| **Primer name** | **Sequence** |
|---|---|
| PP4C-BamHI-F | 5'-GGGGATCCATGGCGGAGATCAGCGACCTGGAC |
| PP4C-EcoRI-R | 5'-GGGAATTCTCACAGGAAGTAGTCGGCCAC |
| PP4R2-EcoRI-F | 5'-GGGAATTCATGGACGTCGAGAGGCTCCAGGAG |
| PP4R2-NotI-R | 5'-<br>TCGCGGCCGCGTCTTGTTCCATTGGTTCATCGGTG |
| PP4R3α-EcoRI-F | 5'-GGGAATTCATGACCGACACCCGGCGGCGG |
| PP4R3α-NotI-R | 5'-GGGAATTCTTATGAATCAAATTTTGCTTT |

| **Vector Type** | **Antibiotic resistance** | **Promotor** | **Fusion Tag** |
|---|---|---|---|
| PcDNA4T0 | Ampicillin | CMV | Flag, HA |
| PCNA3 | Ampicillin | CMV | Flag |

**Table M1. Primers and vectors used for cloning**

#### Silver staining

Proteins in the gels were visualized by silver staining (Nesterenko et al., 1994). All incubations were performed on a shaker at room temperature employing the following steps: The SDS-PAGE gels were first fixed in fixer solution A for 1h and 0,5h in fixer B. After fixing, the gel was sensitized with glutaraldehyde enhancer solution for 1h and washed with distilled water for 2h with several changes of water. After washing, the gel was stained with staining solution for 0,5h. Before developing, the gel was washed three times with distilled water for 10min. The bands were developed with developing solution. The development was stopped with stop solution once the bands were clearly visible.

| **Solution** | **Composition** | **Incubation** |
|---|---|---|
| Fixer A | 50% MetOH 10% Acetic acid | 1h |
| Fixer B | 10%MetOH 5% Acetic acid | 0,5h |
| Enhancer | 30% Gluteraldehyde 25% | 1h |
| Staining | 2g $AgNO_3$, 0,74ml 5N NaOH in 200ml $H_2O$ | 0,5h |
| Developing | 2,5ml Nitric acid 0,25ml Formaldehyde in 500ml $H_2O$ | 5-10min |
| Stop | 50% MetOH 10% Acetic acid | 1h |

**Table M2: Silver staining**

#### Plasmid Transfection

One day before transfection, cells were plated in 100-mm dishes at 40-50% confluence. After one-day growth, transfections were performed with Polyethyleneimine (PEI, Sigma, St. Louis, MO, USA) using 4µL PEI (stock, 1mg/mL) per µg of DNA in serum free medium and incubated for 5 min at room temperature. The mix was added to the cell culture and allowed to grow at 37ºC, in the incubator humidified 5% $CO_2$ atmosphere for 48 hours.

#### Polymerase chain reactions for Site-directed Mutagenesis.

PCR reactions were performed using 5-15 ng of templates DNA in 25µL of GoTaq mix, 0.3µM of each primer and H20 (Milli-Q) added up to 50µl. The DNA was amplified during 35 cycles of denaturation at 95ºC for 30 seconds; annealing temperature was approximately 5ºC lower than calculated $T_m$ of the primers, extension at 72ºC for 60 seconds per 1 kb. The final extension was performed at 72ºC for 7 minutes and final hold, 4°C. The non-mutated DNA was digested with 3U of *Dpn*I restriction enzyme for 1h at 37ºC. Following, the reactions was transformed and positive clones selected. PCR products were verified by sequencing.

| **Primer name** | **Forward Primer (5' to 3')** | **Reverse Primer (5' to 3')** |
|---|---|---|

| | | |
|---|---|---|
| PP4R3A K64R | CTAACACTGCATACCAGAGACAA CAGGACACTCTG | CAGAGTGTCCTGTTGTCTCTGGT ATGCAGTGTTAG |
| PP4R3A K64Q | GCA TAC CAG CAA CAA CAG GAC ACT CTG | GTC CTG TTG TTG CTG GTA TGC AGT GTT AGG |
| PP4R3B K64R | CAAATACTGCATATCAGAGACAA CAGGATACATTAATTG | CAATTAATGTATCCTGTTGTCTC TGATATGCAGTATTTG |
| PP4R3A K64Q | GCA TAC CAG CAA CAA CAG GAC ACT CTG | GTC CTG TTG TTG CTG GTA TGC AGT GTT AGG |
| PP4R3A K642R | GAAAGGCAAGATAATCCCAGACT TGACAGTATGCG | CGCATACTGTCAAGTCTGGGATT ATCTTGCCTTTC |
| PP4R3A 2K64Q | CATACCAGCAACAACAGGAAGAT GAAAAATTTC | CTTCCTGTTGTTGCTGGTATGCA GTGTTAGG |
| PP4R3B K806R | CTAATGGATCCTCTTCCAGAACC ACAAACTTGCC | GGCAAGTTTGTGGTTCTGGAAG AGGATCCATTAG |
| PP4R3B K806Q | CTAATGGATCCTCTTCCCAAACC ACAAACTTGCC | GGCAAGTTTGTGGTTTGGGAAG AGGATCCATTAG |
| PP4R3A D67N | GCATACCAG AAACAACAGAACACTCTGATTG | CAGACCACACAATCAGAGTGTTC TGTTGTTTC |

**Table M3. Primers used for mutagenesis**

**Whole cell extract.**

Cells were harvested using a *cell scraper*, nuclear and cytoplasmatic extracts were prepared as previously described by Dignam (Dignam et al., 1983). Briefly, cells were washed two times with PBS and lysed in buffer A (10 mM Tris, pH 7.8, 1,5 mM MgCl2, 10 mM KCl, 0,1 mM PMSF, 0,5 mM DTT, protease inhibitor cocktail (sigma)) for 10 minutes on ice, centrifuged at 12000 x g for 1 minute at 4°C. The supernatant (the cytoplasmatic fraction) was transferred into a clean tube. The nuclear fraction was lysed in buffer C (10 mM Tris, pH 7.8, 1,5 mM MgCl2, 0.42 mM NaCl, 25% glycerol, 0.2% EDTA, 0,1 mM PMSF, 0,5 mM DTT, protease inhibitor cocktail (sigma)) during 20 minutes on ice. The nuclear fraction was harvested by centrifuging the lysate at 12,000 x g for 3 minutes at 4°C, used or stored separately at -80ºC until further use.

**Co-immunoprecipitation (CoIP)**

For immunoprecipitation experiments, cell extracts were incubated with either α-FLAG, or α-HA resin (Sigma-Aldrich) overnight. Beads were washed three times with BC100 buffer (10mM Tris pH 7.8, 0.5 mM EDTA, 0.1mM PMSF, 0.1 mM DTT, 10% glycerol, 100 mM KCl) and 3 times with BC500 buffer (500 mM KCl). Thereafter, the proteins were eluted with 0.2 M Glycine pH 2 or by incubation with the corresponding competing peptides for enzymatic assay or mass spectrometry analysis

**SDS-PAGE and western blot analysis**

Protein samples were mixed with 5x Laemmli sample buffer (supplemented with 10% *β* -Mercaptoethanol) and boiled at 95ºC for 2 minutes before loading. Protein extracts were fractionated by SDS-PAGE (10% polyacrylamide) and then transferred to nitrocellulose membrane in in transfer buffer (500 mM glycine, 50 mM Tris-HCl, 0.01% SDS, 20% methanol). The membrane was blocked with 5% (w/v) nonfat milk (in Tris-buffered saline, containing 0.01% Triton-X-100), and then the membrane was incubated with primary Antibodies for 1h to overnight at 4℃, After washing the membranes three times for 5 minutes each, followed by incubation with appropriate secondary for 30 minutes. After three times for 5 minutes, antibody binding was visualized with an ECL chemiluminescence system (Millipore detection kit) and short exposure of the membrane to X-ray films or iBright® Imaging Systems (Thermo Fisher Scientific).

**Detecting deacetylation of PP4R3A/B (K64)**

Biotinylated peptides (Biotin-PNTAYQK(Ac)QQDTLI) previously immobilized on magnetic Streptavidin beads (Dynabeads MyOne Invitrogen™ 65601) was incubated with either 1) $H_20$ , 2)$NAD^+$ (0.5 nM) or 3) SIRT1 and $NAD^+$ (0.5 nM) in a total volume of 100 µl of deacetylase buffer (60 mM Tris HCl pH 7,8, 40 mM MgCl2, at pH 7.0, 2mM DTT and protease inhibitor cocktail).The reaction mixtures were incubated at 37 °C for 90 minutes and the beads were washed three times with BC100 and deacetylation of Ac- PP4R3A/B (K64) by SIRT1 was detected by mass spectroscopy .

| | 1 | 2 | 3 |
|---|---|---|---|
| Deacetylation buffer(10X) | 10 µl | 10 µl | 10 µl |
| Beads + peptide | 50 µl | 50 µl | 50 µl |
| $NAD^+$ (Stock 50nM) | - | - | 5 µl |
| SIRT1 | - | 30 µl | 30 µl |
| $H_20$ | 40ul | 10 µl | 5µl |

**Table M4.** Biotinylation reactions

**Immunofluorescence assay (IF) and high-throughput microscopy (HTM)**

The cells were placed on glass coverslips in a 6-well plate, one day before treatment and immunofluorescence assay was performed as follows. Cells were washed two times with PBS (pH 7.4) and fixed with freshly prepared 4% paraformaldehyde (in PBS) for 7 minutes at room temperature (RT). In the case of SIRT1 MEFs, before fixation, the cells were treated with pre-extraction buffer (25 mM Hepes, pH 7.4, 50 mM NaCl, 1 mM EDTA, 3 mM $MgCl_2$, 300 mM sucrose, and 0.5% Triton X-100) for 10 minutes on ice. Fixed cells were then washed two times for 5 minutes in cold PBS and permeabilized in Buffer B (3% BSA, 0.2% Triton-X-100 in PBS) and incubated with Blocking buffer (3% BSA diluted in PBS) for 1 hour at room temperature. Thereafter, the cells were washed three times in blocking buffer and incubated with appropriate primary antibody overnight. Subsequently, the cells were washed three times in blocking buffer and incubated with appropriate secondary antibody conjugated to Alexa Fluor dyes, 488 (green), 594(Texas Red), or 647 (red) (Invitrogen) for 30 minutes at RT in the dark. For DNA staining, DAPI stain was used (1:10000 in PBS1X). DAPI-stained cells were incubated for 5 minutes at

RT after last wash of secondary antibody, followed by two more washes with PBS. After staining, coverslips were mounted in Mowiol and allowed to dry overnight before imaging. Images were viewed and captured on a Leica SP5 microscope (Leica, Milton Keynes, UK). Primary antibodies and dilutions were: RPA32 (Cell signalling) at 1:150, anti-Phospho RPA32(Novus) and anti-Phospho RPA32 (S4/S8) (Sigma-Aldrich) at 1:200, Phospho-yH2AX (Abcam) at 1: 500. In the case of high-throughput microscopy (HTM) immunofluorescence, SIRT1 MEFs were grown in LabTek II Chamber slides (Nunc, Thermo Fisher Scientific) and stained using the procedure mentioned above. For HTM, 48 images were automatically acquired per well with a Scan^R (Olympus) with an oil immersion objective at × 40 magnification and non-saturating conditions. Automated image segmentation of DAPI-stained nuclei was generated from which the corresponding signals were calculated using Fiji Software (https://fiji.sc/) and a package based on the Cell Profiler.

Confocal fluorescence images were obtained on a Leica DM2500 SPE confocal system. Images were taken with 40x NA 1.15 oil or 63x NA 1.3 oil objectives and the standard LAS-AF software. Possible crosstalk between fluorochromes was avoided by carefully adjusting laser intensities and HyD gain, thus avoiding false-positive colocalization signals. For high-throughput microscopy (HTM), 24-48 images were automatically acquired from each well with a robotized fluorescence microscopy station (ScanR; Olympus) at 40x magnification and non-saturating conditions. Images were segmented using the DAPI staining to generate masks matching cell nuclei from which the corresponding signals were calculated using an in-house–developed package based on Cell Profiler (www.cellprofiler.org), an open-source software program. In collaboration with the Advanced Digital Microscopy Facility at IRB Barcelona, we developed a pipeline to load the stack of 8-bit images with 3 channels, generate nuclear masks with the DAPI channel and measure mean intensity of the two additional channels. Nuclei had a typical diameter of 60–150-pixel units, and background fluorescence in DAPI images below an absolute threshold of 0.20-0.35 was set to 0. These results were exported to Excel for further analysis and GraphPad-Prism was used for graphical representation.

**Generation of gRNAs and CRISPR/Cas9 gene editing**

The gRNAs designed sequences for SIRT1 are as follows: gRNA-1: acctttgcaaacttgaagaa, gRNA-2: aacaggttgcgggaatccaa, gRNA-3: gttgactgtgaagctgtacg. They were cloned into pSpCas9 (BB)-2A-GFP (PX458) (https://www.addgene.org/4813) according to published protocol (Ran et al., 2013)and the resulting constructs were sequenced to verify successful gRNA integration. The U2OS cells were transfected with all three constructs using PEI and were sorted for GPF positive cells 48h after transfection. Single clones of the cells were produced using serial dilution method and were screened for SIRT1 knockout by WB and PCR.

**siRNA transient interference assay**

PPP4C siGENOME SMARTpool siRNA was purchased from Dharmacon (Lafayette, CO) (Cat. #D-008486-01, D-008486-03, D-008486-04, D-008486-05). For transient transfection, the U2-OS cells were seeded 5×10$^3$/ml on a 6-well plate with antibioticsfree medium. Following incubation overnight, targeting siRNA was transfected with siPPP4C or the same amount of scrambled control siRNA (Cat. # D-001206-13-05 non-targeting pool) using DharmaFECT1 transfection reagent (Dharmacon) according to the manufacturer's instructions. Following incubation for 48 h, the cells were harvested or treated with EdU for the indicated period of time.

### Mass spectrometric analysis

SIRT1-PP4 identification

SDS-PAGE resolved and gel bound protein bands shown in Figure 1 were subjected to in-gel digestion followed by peptide clean up using 2µL bed-volume of Poros 50 R2 reversed-phase beads packed in Eppendorf gel-loading tip (RP-microtip column), following published protocol (Erdjument-Bromage et al., 1998). Matrix Assisted Lasers Desorption Reflectron Time of Flight Mass spectrometry (MALDI-ReTOF MS) was carried out on two peptide pools each (16% and 30% acetonitrile) recovered from the RP-microtip column using a Bruker Reflex III instrument with delayed extraction. Three internal peptide calibrants (synthetic peptides) of known molecular weights were used to internally calibrate the peptide pools obtained from the samples. For mass fingerprinting protein identification, top 'major' experimental masses (m/z) combined from MALDI-ReTOF experiments were used to search human (20,380 entries, June 2018) database using Mascot proteomics software (Version 2.5.0, Matrix Science, London, UK. www.matrixscience.com ) A mass accuracy better than 50ppm and maximum two missed cleavage site was allowed per peptide. A detailed Table (Table S1) with top identified protein details such as protein sequence coverage, peptide spectral matches, can be found.

Identification of acetylated peptides in PP4 complex in Wt and SIRT1 KO cells

PP4 complex was purified from MEFs *Wt* and *Sirt1*$^{-/-}$ overexpressing PP4 core complex (PP4c and PP4R2). The samples were precipitated overnight with cold acetone to remove the glycerol and other reagents can interfere with the protein digestion. Briefly, after samples precipitation the samples were diluted in 50 µL of 8 M urea/ 50 mM NH4HCO3(pH 8.5) solution by ultrasonic bath and mixing. Proteins were reduced (DTT 20 mM50 in mM NH4HCO3; 90minutes, 32ºC) and alkylated (iodoacetamide 35 mM in 50 mM NH4HCO3; room temperature for 30 min, in the dark). Afterwards, the samples were diluted down to 1 M urea with digestion buffer and proteins were digested with trypsin (Sequence grade modified porcine Trypsin, Promega; 1µg trypsin/sample, pH 8, 34oC, 16h overnight). The resulting peptide mixtures were cleaned-up with C18 tips (PolyLC Inc.) as per manufacturer´s protocol. Finally, the cleaned-up peptide solutions were dried-down and stored at -20ºC until the analysis. The peptide samples (Table S2) were analyzed using a LTQ-Orbitrap Velos, mass spectrometer (Thermo Fisher Scientific, San Jose, CA, USA) coupled to an nanoAcquity liquid chromatographer (Waters). The tryptic digests were resuspended in 1% FA solution and an aliquot of each was injected for chromatographic separation. Peptides were trapped on a Symmetry C18TM trap column (5µm180µm x 20mm; Waters), and were separated using a C18 reverse phase capillary column (ACQUITY UPLC BEH column; 130Å, 1.7µm, 75 µm x250mm, Waters). The gradient used for the elution of the peptides was 1 to 40 % B in

90 minutes, followed by gradient from 40% to 60% in 10 min (A: 0.1% FA; B: 100% ACN, 0.1%FA), with a 250 nL/min flow rate.
Eluted peptides were subjected to electrospray ionization in an emitter needle (PicoTip$^{TM}$, New Objective) with an applied voltage of 2000V. Peptide masses (*m/z* 300-1700) were analyzed in data dependent mode, where a full Scan MS was acquired in the Orbitrap with a resolution of 60,000 FWHM at 400m/z. Up to the 15$^{th}$ most abundant peptides (minimum intensity of 500 counts) were selected from each MS scan and then fragmented in the linear ion trap using CID (38% normalized collision energy) with helium as the collision gas. The scan time settings were: Full MS: 250 ms (1 microscan) and MSn: 120 ms. Generated. raw data files were collected with *ThermoXcalibur(v.2.2)*.
Searches were performed with the. raw data obtained in the mass spectrometry analyses using Thermo Proteome Discover (v.1.4.1.14) with SequestHTas the search engine against SwissProt Human and mouse public Database (version of January 2017).
The following search parameters were applied trypsin enzyme with 2 missed cleavages, carbamidomethyl of cysteine as a fixed modification and acetylation and oxidation of methionine as a variable, the peptide tolerance 10ppm and 0.6 Da for MS and MS/MS spectra. To improve the sensitivity of the database search, Percolator (semi-supervised machine learning) was used in order to discriminate correct from incorrect peptide spectrum matches. Percolator assigns a q-value to each spectrum, which is defined as the minimal FDR at which the identification is deemed correct. These q-values are estimated using the distribution of scores from the decoy database search. The results have been filtered, so only proteins identified with at least 2 high confidence peptides (FDR≤ 0.01, validation based on q-value) are included in the table S2.

<u>K64 deacetylation by SIRT1</u>

In order to evaluate the role of SIRT1 in deacetylation we performed an *in vitro* reaction with SIRT1 in the presence or absence of the cofactor NAD+ using a biotin-PNTAYQK(Ac)QQDTLI peptide bound to Streptavidin magnetic beads as substrate.
After the reaction, the beads were washed four times with PBS and subsequently eluted with elution buffer (80% acetonitrile, 0.2% trifluoracetic acid and 0.1% formic acid). Additionally, an aliquot of pure peptide was cleaned-up using a C18 reverse phase column before to the analysis. Both, the eluted and the pure peptide were mixed at 1:1 ratio with CHCA (α-cyano-4-hydroxycinnamic acid), spotted into an Anchorchip plate and left to dry. The mass spectra were acquired using AutoFlex MALDI-TOF mass spectrometer (Bruker Daltonik, Bremen, Germany), reflector type time-of-flight mass spectrometer, equipped with a pulsed nitrogen laser working at 337 nm and a smartbeam II laser working at 355 nm, respectively. The Autoflex instrument is operated in the positive ion mode with delayed extraction at an accelerating voltage of 20 kV and a variable voltage reflectron.
